# The role of wingbeat frequency and amplitude in flight power

**DOI:** 10.1101/2022.06.28.497935

**Authors:** Krishnamoorthy Krishnan, Baptiste Garde, Ashley Bennison, Nik C. Cole, Emma-L. Cole, Jamie Darby, Kyle H. Elliott, Adam Fell, Agustina Gómez-Laich, Sophie de Grissac, Mark Jessopp, Emmanouil Lempidakis, Yuichi Mizutani, Aurélien Prudor, Michael Quetting, Flavio Quintana, Hermina Robotka, Alexandre Roulin, Peter G. Ryan, Kim Schalcher, Stefan Schoombie, Vikash Tatayah, Fred Tremblay, Henri Weimerskirch, Shannon Whelan, Martin Wikelski, Ken Yoda, Anders Hedenström, Emily L.C. Shepard

## Abstract

Body-mounted accelerometers provide a new prospect for estimating power use in flying birds, as the signal varies with the two major kinematic determinants of aerodynamic power: wingbeat frequency and amplitude. Yet wingbeat frequency is sometimes used as a proxy for power output in isolation. There is therefore a need to understand which kinematic parameter birds vary and whether this is predicted by flight mode (e.g., accelerating, ascending/descending flight), speed or morphology. We investigate this using high-frequency acceleration data from (i) 14 species flying in the wild, (ii) two species flying in controlled conditions in a wind tunnel and (iii) a review of experimental and field studies. While wingbeat frequency and amplitude were positively correlated, R^2^ values were generally low, supporting the idea that parameters can vary independently. Indeed, birds were more likely to modulate wingbeat amplitude for more energy-demanding flight modes, including climbing and take-off. Nonetheless, the striking variability even within species and flight types, highlights the complexity of describing the kinematic relationships, which appear sensitive to both the biological and physical context. Notwithstanding this acceleration metrics that incorporate both kinematic parameters should be more robust proxies for power than wingbeat frequency alone.

## Introduction

Factors affecting the energetic costs of flight can have a profound influence on the ecology and behaviour of birds, with flight conditions affecting the location of migratory flyways, and in particular cases, breeding success (Kranstauber et al., 2015; Weimerskirch et al., 2012). Yet at fine-scales, disentangling the impact of the biological and physical environment on flight costs can be challenging, given that a range of factors often vary simultaneously. These include the topography birds are flying over, individual position within a flock (Garde et al., 2021, Portugal et al., 2014) and social context (Sankey & Portugal, 2019; Usherwood et al., 2011), as well as factors that vary over longer timescales including the birds’ immunological state (Hicks et al., 2018), and physical factors such as wind speed, turbulence and air density (Bishop et al., 2015; Furness & Bryant, 1996; Sapir et al., 2010). High-frequency data from animal-attached loggers have proved powerful in this regard, as the signal from onboard accelerometers can be used to quantify second-by-second changes in wingbeat frequency (Cochran et al., 2008; Sato et al., 2008; Van Walsum et al., 2020), and potentially other kinematic parameters (Taylor et al., 2019).

Power varies in a U-shaped fashion with flight (air) speed for most flying birds (Engel et al., 2010; Hedenström & Lindström, 2017; Norberg, 2012; Pennycuick, 2008; Tobalske et al., 2003), and wingbeat frequency seems to follow the same trend, although it is not always pronounced (Ellerby & Askew, 2007; Hedrick et al. 2003; Pennycuick et al., 1996; Schmidt-Wellenburg et al., 2007; Tobalske et al., 2003; Usherwood et al., 2011). This explains why wingbeat frequency has been used as a proxy for flight costs in a range of ecological studies (e.g., Taylor et al., 2019; Usherwood et al., 2011). However, wingbeat frequency also has limitations as a proxy for power requirements, because studies by Hedrick et al. (2003) and Tobalske et al. (2003) have shown that the minimum wingbeat frequency does not always coincide with the minimum power speed. In fact, it can occur at over twice the minimum power speed, which demonstrates that other kinematic parameters, such as wingbeat amplitude, stroke plane angle, and span-ratio can have an important role in modulating power output (Pennycuick et al., 2000; Rosén et al., 2004, 2007; Ward et al., 2001).

The major determinants of the aerodynamic power output of a flapping wing are the wingbeat frequency (*f*) and amplitude (*A*). In flapping flight, the resultant aerodynamic forces (lift, drag and thrust) acting on the wing are predominantly determined by the flow over each wing section at each time instant (Shyy et al., 2010). This is the combination of the flow due to the forward motion (forward velocity) of the bird and the flapping motion of the wing (wing velocity). The flow over the wing section can be controlled by the wing velocity, which solely depends on the wingbeat frequency and wingbeat amplitude (Pennycuick, 2008). The aerodynamic forces exerted on the wings are proportional to the square of the velocity, and the mechanical power output is proportional to the cube of the velocity (Pennycuick, 2008). Therefore, while the total resultant aerodynamic forces can be modulated by varying the wing planform and angle of attack during flight, modulating the flow velocity over each wing section has the major effect. The power can be shown to be proportional to the cube of both amplitude and frequency, if the product of wingbeat amplitude and frequency is substituted for velocity (as they both scale the same with velocity (Floryan et al., 2018)):

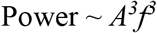

Despite the importance of both wingbeat frequency and amplitude for overall power output, an overview of the scenarios under which birds modulate one or the other parameter is lacking. Indeed, examples from the literature suggest that the relationship is not straightforward: Some studies show that birds vary their power output with little to no change in wingbeat frequency (Tobalske & Biewener, 2008; Torre-Bueno & Larochelle, 1978; Wang et al., 2019), whereas others report that wingbeat frequency varies with the power output while the amplitude is unaltered (Ellerby & Askew, 2007). It is therefore unclear whether birds vary frequency or amplitude to modulate power according to their flight mode (e.g., hovering, climbing, manoeuvring, or level flight) or morphology.

Power can theoretically be modulated either by a contribution from both wingbeat frequency and amplitude, or by changes to one or the other. What is clear is that a proxy for flight power should ideally integrate information on wingbeat frequency and amplitude in order to be widely applicable. Two related proxies for energy expenditure have been proposed using data from body-mounted accelerometers, both of which integrate information on stroke frequency and signal amplitude. Dynamic Body Acceleration (DBA) was proposed in 2006 as a metric that captures whole-body acceleration (Wilson et al., 2020, 2006), and has been shown to vary with the energy expended by free-living auks (Elliott et al., 2013) and cormorants (Hicks et al., 2017) in flight. However, the precise relationship between the DBA signal and wingbeat kinematics is unknown. Spivey and Bishop (2013) also established a theoretical framework of how body acceleration can be related to the biomechanical power output of flapping flight, using the root mean square values of heave and surge acceleration and wingbeat frequency.

This assumes that the amplitude of the dorsoventral or “heave” accelerometer measurements vary with the wingbeat amplitude (Usherwood et al., 2011). However, similar to DBA, the relationship between body and wing motions, and how they covary over a wingbeat cycle, has not been established.

In this study, we examine the outlook for acceleration-based proxies for power use in flapping flight across species and contexts. Specifically, we (1) test how the output of body-mounted accelerometers varies with wingbeat amplitude, using a novel methodology, (2) assess whether birds preferentially use wingbeat frequency or amplitude to modulate their power output (or a proxy such as speed) according to (a) their body mass or morphology, and (b) their flight mode. We address this by reviewing the experimental literature, where wingbeat kinematics have largely been quantified using high-speed video, and by conducting further trials, where we equip 14 species of bird with body-mounted accelerometers to monitor their flight behaviour in the wild.

## Methods

### (i) Wind tunnel trials: Does the acceleration signal vary with wingbeat amplitude?

Movement of the wings results in movement of the body in the same axis. Greater wingbeat amplitudes should result in greater vertical accelerations of the body for a fixed wingbeat frequency. We examined these relationships using a body-mounted accelerometer and magnetometer, and a small neodymium boron magnet attached to the leading edge of the wing (Wilson & Liebsch, 2003). The geomagnetic signal strength in each axis varied throughout the wingbeat cycle as a function of the angle and distance to the magnet. We therefore calculated the vector sum from all three magnetometer channels, which varied solely with the distance to the magnet, giving a clear peak per wingbeat cycle when the magnet was closest to the sensor. This allowed us to assess how the vertical body acceleration varied in relation to the maximum vector sum from the magnetometer (as a proxy for wingbeat amplitude) within the same wingbeat cycle.

Data were collected from two species flying at a range of speeds in large, low turbulence wind tunnels. In one set of trials, two pigeons (*Columba livia*) were equipped with Daily Diary (DD) data loggers (Wildbyte Technologies, Swansea University, UK), sampling acceleration at 150 Hz and magnetic field strength at 13 Hz. Each pigeon was equipped with two units; one on the upper back and another on the lower back. The logger at the top of the back was positioned close to the magnet, whereas the logger on the lower back was sufficiently far from the magnet not to be influenced by it (as determined in preliminary tests). The second logger allowed us to control for the potential influence of changing geomagnetic field strength (due to changes in bird trajectory) on the magnetometer output. Loggers had dimensions of 22 × 15 × 9 mm and a total mass (3.4 g per logger and battery) that was less than 3% of the bird body mass. A cylindrical neodymium boron magnet (8 × 2 mm, 0.19 g) was taped to the leading edge of the wing, close to the wing root (Figure 1 A). Both the loggers and the magnet were attached with micropore tape. Pigeons were flown at speeds between 12 and 18 m·s^-1^. Experiments were performed between 25/01/2019 and 01/02/2019 in the wind tunnel of the Max Planck Institute for Ornithology, Germany, under ethical approval Gz.: 55.2-1-54-2532-86-2015 granted by the government of Upper Bavaria (Sachgebiet 54 – Verbraucherschutz, Veterinärwesen, 80538 München), and Swansea University AWERB, permit number 030718/66.

**Figure 1.**
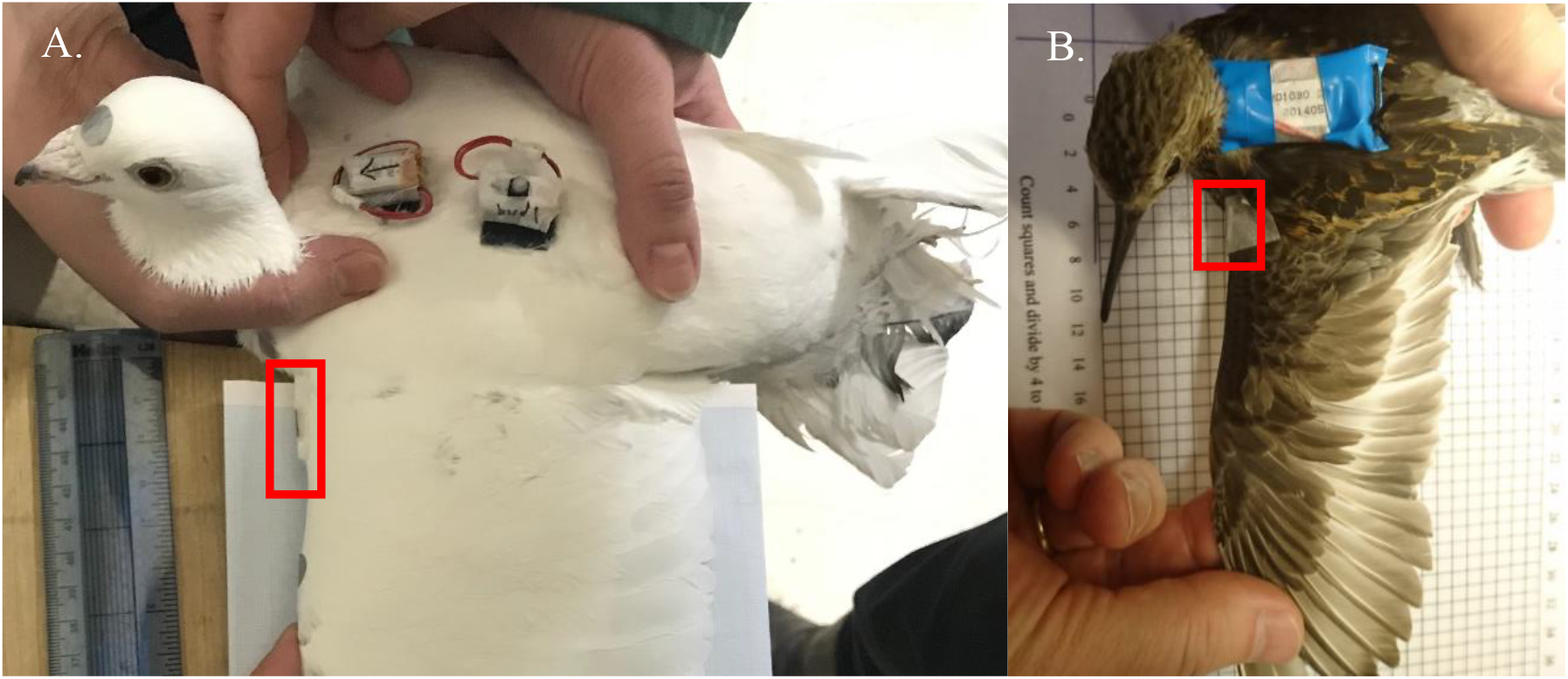
Setup of the tag (DD; containing both an accelerometer and magnetometer) and magnet (highlighted by the red rectangle) on A. a pigeon and B. a dunlin.

Further trials were conducted with a dunlin (*Calidris alpina*) in the wind tunnel at Lund University, Sweden, which has similar performance characteristics to the tunnel in Seewiesen (Pennycuick et al., 1997). A small neodymium magnet (4 × 2 mm, 0.02 g) was attached to the wing of the dunlin following the same procedure. A single unit logging tri-axial acceleration and magnetic field strength at 100 Hz (Technosmart Europe) was attached to the back of the dunlin with a backpack harness (Figure 1 B). The logger was 16 × 24 × 12 mm and weighed 2.6 g, equivalent to 4.8% of the bird’s body mass. The dunlin was flown at a range of speeds for < 10 minutes. Ethical permission was obtained from Swansea University AWERB, permit number 030718/66.

### (ii) Variation in the amplitude - frequency relationship across species

Data from birds flying in the wind tunnel were combined with acceleration data from a further 12 species of free-flying birds (Table 1) to examine relationships between wingbeat frequency and amplitude, and whether birds are more likely to use one parameter or the other to modulate their power output, according to their mass and morphology. Datasets were selected for inclusion according to whether tags were attached on the back, rather than the tail, to minimize the contribution of the angular motion of the bird to the acceleration signal, when the sensor is placed far from the centre of mass (Garde et al., 2022).

**Table 1.** Datasets in the study, along with the number of individuals tracked, body mass, wingspan and wing area, and the source of the morphometric data.

| Species | Location | N | Mass (g) | Wing span (m) | Wing area (m<sup>2</sup>) | Tag type | Data from literature | Source |
| --- | --- | --- | --- | --- | --- | --- | --- | --- |
| **Brünnich's guillemot**<br>*Uria lomvia* | Coats Island, Nunavut, Canada | 13 | 949 | 0.727 | 0.069 | Daily<br>Diary | Wings | Orben et al., 2015 |
| **Common guillemot**<br>*Uria aalge* | Puffin Island, UK | 6 | 1050 | 0.73 | 0.056 | AxyTrek | Mass,<br>Wings | Spear & Ainley, 1997 |
| **Northern fulmar**<br>*Fulmarus glacialis* | Saltee Islands, Ireland | 3 | 778 | 1.12 | 0.106 | Daily<br>Diary | Wings | Warham, 1977 |
| **Pigeon**<br>*Columba livia* | Radolfzell, Germany | 9 | 456 | 0.647 | 0.064 | Daily<br>Diary | None | Measured directly |
| **Red-tailed tropicbird**<br>*Phaethon rubricauda* | Round Island, Mauritius | 10 | 820 | 1.115 | 0.117 | Daily<br>Diary | None | Measured directly |
| **Great frigatebird**<br>*Fregata minor* | Europa Island | 3 | 1113 | 2.084 | 0.365 | Daily<br>Diary | None | Measured directly |
| **Black-legged kittiwake**<br>*Rissa tridactyla* | Middleton Island, Alaska, USA | 3 | 387 | 0.965 | 0.101 | Daily<br>Diary | Wings | Pennycuick, 1997 (n = 2) |
| **Imperial cormorant**<br>*Leucocarbo atriceps* | Punta Leon, Argentina | 5 | 2400 | 1.13 | 0.183 | Daily<br>Diary | Mass,<br>Wings | Quintana et al., 2011, Spear & Ainley, 1997 (n = 1) |
| **Western barn owl**<br>*Tyto alba* | Switzerland | 10 | 296 | 0.936 | 0.134 | AxyTrek | None | Measured directly |
| **Grey-headed albatross**<br>*Thalassarche chrysostoma* | Marion Island,<br>South Africa | 5 | 3290 | 2.186 | 0.348 | Daily<br>Diary | Mass,<br>Wings | Phillips et al., 2004 (n = 1) |
| **Wandering albatross**<br>*Diomedea exulans* | Marion Island,<br>South Africa | 6 | 8500 | 3.01 | 0.583 | Daily<br>Diary | Mass,<br>Wings | Pennycuick, 1997;<br>Pennycuick, 2008 |
| **Streaked shearwater**<br>*Calonectris leucomelas* | Awashima Island,<br>Japan | 5 | 503 | 1.119 | 0.126 | Daily<br>Diary | Wings | Shirai et al., 2013 |
| **Dunlin**<br>*Calidris alpina* | Sweden | 1 | 55 | 0.334 | 0.014 | Axy XS | Mass,<br>Wings | Hentze, 2012 |
| **Northern gannet**<br>*Morus bassanus* | Saltee Islands,<br>Ireland | 10 | 2856 | 1.85 | 0.262 | Axy | Wings | Spear & Ainley, 1997 (n = 1) |

Morphological parameters including wing loading, wingspan, wing area, and body mass, were either measured directly and averaged (following Pennycuick, 2008) or taken from the literature (Table 1). We used wingspan rather than aspect ratio because there is a framework linking the former to wingbeat kinematics (Pennycuick, 2008). In order to assess the role of wing loading independently from body mass, we calculated the residuals of the linear regression between log(wing loading) and log(body mass) (Lee et al., 2008).

All birds flying in the wild were equipped with tags recording tri-axial acceleration at 40 Hz (except common guillemots and gannets, where the sampling rate was 50 Hz and pigeons, where it was 180 Hz). An examination of accelerometer data revealed some slight variation in sampling rate between logger types (up to 3 Hz), which was accounted for in the calculation of wingbeat frequency. Tags were attached to the back feathers using Tesa tape (Wilson et al., 1997) in all species apart from pigeons, where tags were attached via Velcro strips glued to the back feathers (Garde et al., 2021, Biro et al., 2002). The total mass of the tag, including housing and attachments, was under 5% of bird body mass and 3% in most cases. See Table SI 1 for details of ethical permissions.

Episodes of flapping flight were identified visually from the acceleration data (Shepard et al., 2008). Only periods of consistent flapping, with no interruption or rapid changes in amplitude, were selected for the analysis of both wind tunnel and wild data, irrespective of the species. Wingbeat frequency and heave amplitude (amplitude of the vertical body acceleration within a wingbeat) were quantified using the following approach, which enabled the estimation of the period of individual wingbeats. Peaks in heave acceleration associated with the downstroke (Figure 2) were identified by smoothing raw heave values over 3-5 datapoints for all species except the guillemots, which did not require smoothing as their high wingbeat frequency resulted in a relatively clean signal. A second-order derivative was then applied to identify the positive-to-negative turning points. Peaks were marked when the differentials exceeded a threshold within 5 points of the turning point. Thresholds were manually selected for each flight bout so that they only captured wingbeat peaks, as characterised by high heave accelerations (around 2 *g*). The section between each marked peak was considered as one wingbeat cycle and used to determine the wingbeat period (frequency). The wingbeat frequency of the dynamic soaring birds (birds that extract energy by flying through the wind shear in the atmosphere) represents the frequency during the flapping period. The heave amplitude was calculated as the difference between the highest and lowest heave values within the wingbeat. Peak identification was conducted in R version 4.0.2 (Andy Bunn, 2017) using user-defined function for Brunnich’s guillemot, common guillemots, pigeons (homing flights only), and tropicbirds. All other data were processed using custom developed software DDMT (Wildbyte Technologies).

**Figure 2.**
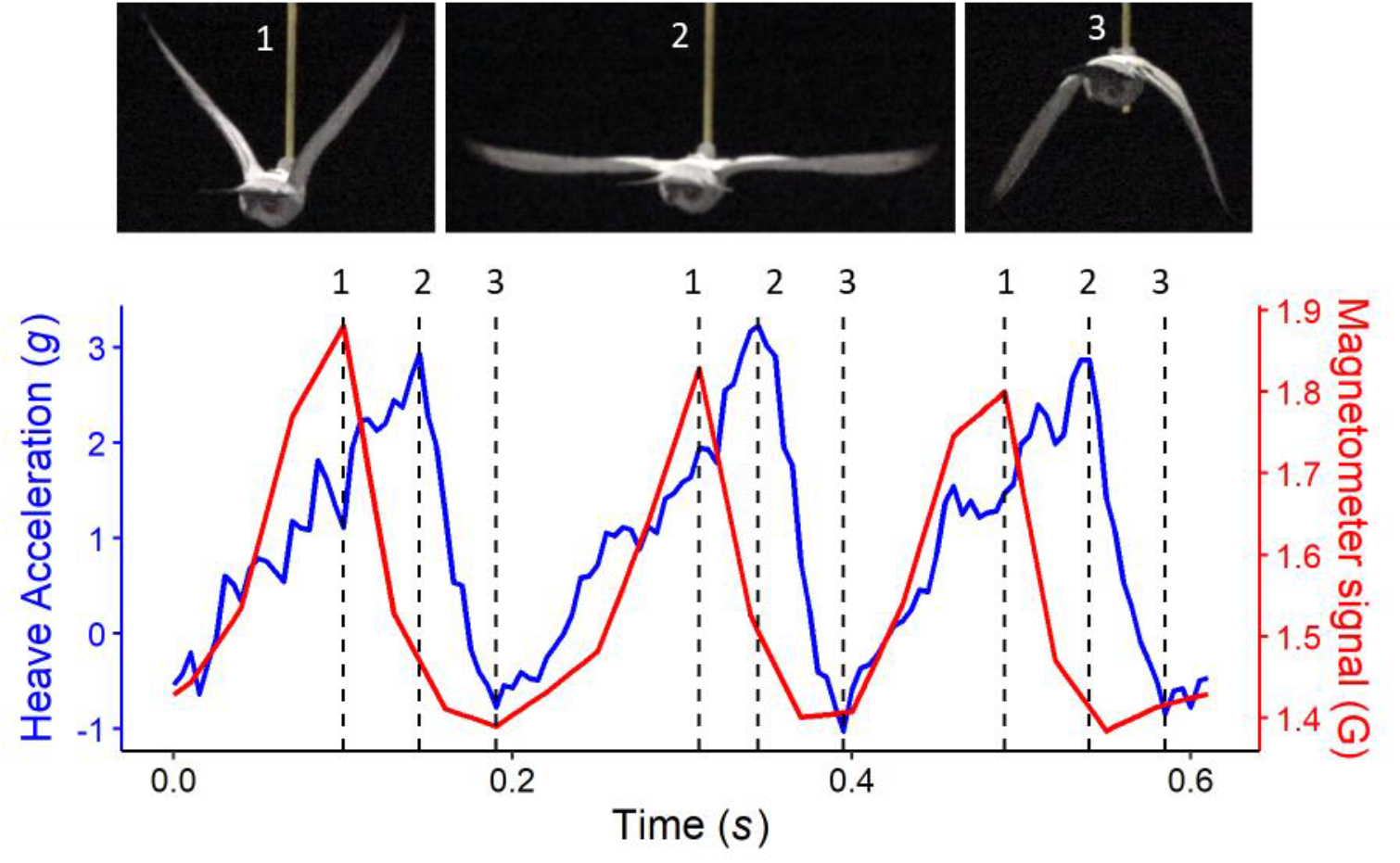
Comparison of the accelerometer (blue) and magnetometer (red) signals in the heave axis for 3 wingbeats from a pigeon flying in a wind tunnel at 15 m s^-1^. (1) Peaks in the magnetometer signal correspond to the start of the downstroke (a smaller acceleration peak is sometimes evident at the same time), (2) peaks in the heave acceleration occur in the middle of the downstroke, and (3) troughs in the magnetometer signal occur at the end of the downstroke. Images from the corresponding wingbeat cycle were captured using a Sony PXW-Z150 camera recording at 120 Hz, which was synchronised with the onboard logger by moving the equipped bird in view of the camera and a clock showing the logger time.

Filters were applied to remove unrealistic wingbeat frequencies. Low outliers were identified during short sections of non-flapping flights that were not excluded during the previous steps. High outliers were also recorded and were probably caused by false peak identification due to rapid manoeuvres. Filtered data were used to estimate final wingbeat frequencies, taken as the average over 10 consecutive wingbeats for wild data (which sometimes occurred in 2 flapping bouts for albatrosses) and 5 wingbeats for wind tunnel data (where the total wingbeats available from consistent flights was lower). Heave amplitude was also averaged over the same interval.

Finally, a simulation confirmed that our ability to estimate signal amplitude across species with variable wingbeat frequencies was not influenced by the sampling frequency (Supplementary information).

### (iii) Variation in wingbeat kinematics with climb rate and airspeed

First, we examined how wingbeat frequency and signal amplitude varied in relation to airspeed for a pigeon flying in the wind tunnel (for which we had reliable records of airspeed). Then we assessed how pigeons, barn owls and tropicbirds varied their wingbeat frequency and amplitude in relation to airspeed and climb rate in the field. These datasets were selected due to the relatively high GPS sampling frequency (1 Hz for pigeons and barn owls, and once per minute for tropicbirds). Airspeed was estimated from the GPS derived groundspeed and the wind vector (Pennycuick, 2008), as recorded by a portable weather station (Kestrel 5500L, Kestrel instruments, USA) mounted on a 5 m pole (see Garde et al., 2021). The weather station was positioned at the pigeons’ release site, and at the highest point of Round Island (280 m a.s.l.) in the case of the tropicbirds. For barn owls, weather data were collected from weather stations located near the owls’ nest. Altitude was calculated from barometric pressure recorded by the Daily Diary (at 4 Hz) in the case of the pigeons and tropicbirds, adjusted for daily changes in sea level pressure (Garde et al., 2021) and climb rate was calculated as the difference between consecutive values of altitude smoothed over 2 s. GPS altitude was used for the barn owls.

Airspeed, climb rate, wingbeat frequency and heave amplitude were averaged over 10 wingbeats for the pigeons and barn owls, and over 1-minute intervals for the tropicbirds (to match the airspeeds). For each interval (10 wingbeats or 1 minute) the proportion of level flapping flight was calculated, and only intervals with ≥ 80% level flapping flight were included in the analysis.

Periods of level flapping flight were selected for the airspeed analysis, taking data where the rate of change of altitude was > -0.2 and < 0.2 m·s^-1^. To minimise the variation in airspeed in the climb rate analysis, we excluded data with airspeeds higher or lower than the overall mean ± 1 standard deviation.

### (iv) Statistical analysis

We used linear models to examine whether the peak heave acceleration increased with the peak magnetometer vectorial sum (as a proxy for wingbeat amplitude) for both dunlin and pigeon wind tunnel flights. We also used linear models to assess whether the heave amplitude varied with wingbeat frequency, using separate models for wind tunnel and wild flights.

To test whether birds varied their wingbeat amplitude to a greater extent than their wingbeat frequency in relation to climb rate and airspeed, we ran separate linear mixed-effects models (LMM) per species (tropicbirds, barn owls and pigeons). These models included wingbeat amplitude as the response variable, expressed as a function of wingbeat frequency and the effect of either airspeed or climb rate on the slope of this relationship (the interaction between wingbeat frequency and either climb rate or airspeed). A positive interaction would indicate that birds increased their amplitude more than frequency to increase speed/ climb rate, while a negative relationship would indicate that they modulate wingbeat frequency more than amplitude. Individual was included as a random factor to account for uncontrolled variation relating to morphology and motivation (only one trip per bird was included). A continuous-time first-order autoregressive correlation structure was included in all models.

To investigate whether morphology affected the degree to which birds varied their wingbeat frequency, we calculated the coefficient of variation for the wingbeat frequency for each species, with the prediction that groups such as auks, with high wing loading, would be constrained in the range of frequencies. We did not run this analysis for the signal amplitude data, as the signal magnitude (and how this varies e.g. with flight speed) might be influenced by factors including device location (Garde *et al*. 2022). We used linear models and Pearson’s product-moment correlation tests to see how the species-specific coefficients of variation (used as response variables) varied with wingspan, body mass, and residual wing loading. Note that pigeon flights recorded in the wind tunnel were not used in this analysis as free flight data had been recorded for pigeons. The dunlin flights were included. All statistical analyses were performed using R version 4.0.2. LMMs were performed using the package “nlme” (Pinheiro et al., 2017, version 3.1-151). Model selection was performed using the package “MuMIn” (Barton & Barton, 2015, version 1.43.17), and the distribution of residuals was tested using “fitdistrplus” (Delignette-Muller et al., 2015, version 1.1-5).

## Results

### (i) Wind tunnel trials: Does the acceleration signal vary with wingbeat amplitude?

Pronounced cyclic changes in the magnetometer signal were evident through the wingbeat cycle for both species that were flown in the wind tunnel (Figure 2) due to the changing magnetic field strength driven by the small magnet attached to the leading edge of the wing. The magnetometer signal was highest at the start of the downstroke, when the distance between the magnet and the transducer was at a minimum, and it decreased as the downstroke progressed, until the magnet was farthest from the logger at the end of the downstroke (Figure 2). In contrast, the maximum heave acceleration occurred mid-downstroke when the wing traversed the body, corresponding to the point of maximal lift generation (Crandell and Tobalske, 2011; Bilo et al., 1984). The magnetometer signal therefore varied with the wing displacement rather than wing (and body) acceleration, explaining why the peaks in magnetic and acceleration signals were offset from each other.

Nonetheless, we found a positive linear relationship between heave amplitude and the peak magnetometer vectorial sum in both species (pigeons: estimate = 1.253, std. error = 1.02, t-value = 5.151, p < 0.001; dunlin: estimate = 2.639, std. error = 0.085, t-value = 31.01, p < 0.001), showing that the body acceleration increases with wingbeat amplitude (Figure 3).

**Figure 3.**
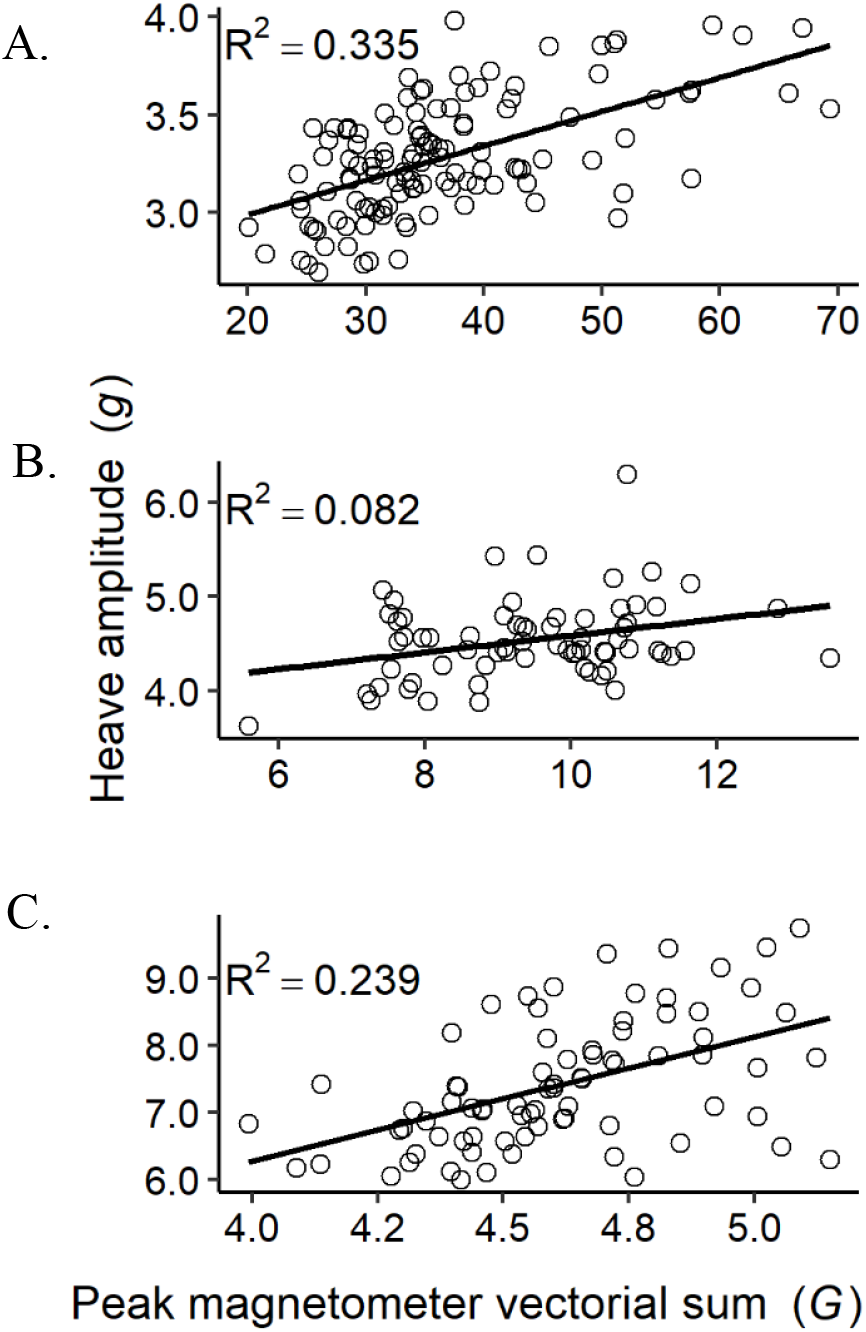
The heave amplitude increased with the maximum magnetometer vectorial sum within wingbeat cycles for A. a dunlin, B. and C. two pigeons flying in wind tunnels across a range of flight speeds. The variation in absolute values from the magnetometer will vary due to the position of the magnet on the wing and its distance to the body-mounted magnetometer. The amplitude of the heave signal is influenced by the position of the back-mounted logger.

### (ii) Assessing the relationship between wingbeat amplitude and frequency

We then assessed how the wingbeat frequency and heave amplitude (as a proxy for wingbeat amplitude as established for pigeon and dunlin) covaried for different species. There was a positive, linear relationship between wingbeat frequency and heave amplitude in almost all species that flew in the wild (n = 13) and the wind tunnel (n = 2) (Table 2). The exceptions were three of the four birds that use dynamic soaring: the northern fulmar, grey-headed albatross, and wandering albatross. Nonetheless, most R^2^ values were relatively low, ranging from 0.001 to 0.38 (Table 2).

**Table 2:** The relationship between heave amplitude and wingbeat frequency for 14 species flying in the wild and 2 species flying in controlled conditions.

| Species | Signal amplitude (g) | Wingbeat frequency (Hz) | Slope | Intercept | p-value | R<sup>2</sup> | Total wingbeats |
| --- | --- | --- | --- | --- | --- | --- | --- |
| Dunlin* | 3.2 ± 0.3 | 13.2 ± 0.9 | 0.110 | 1.833 | < 0.001 | 0.112 | 73 |
| Pigeon* | 6.0 ± 0.7 | 6.7 ± 0.4 | 0.893 | 1.337 | < 0.001 | 0.309 | 147 |
| Pigeon | 3.7 ± 0.4 | 5.2 ± 0.5 | 0.189 | 2.713 | < 0.001 | 0.048 | 4,858 |
| Barn Owl | 2.4 ± 0.4 | 4.4 ± 0.4 | 0.518 | 0.531 | < 0.001 | 0.162 | 134,919 |
| Common Guillemot | 2.5 ± 0.3 | 9.7 ± 0.6 | 0.206 | 0.541 | < 0.001 | 0.170 | 31,349 |
| Brünnich's Guillemot | 1.3 ± 0.2 | 7.7 ± 0.5 | 0.180 | -0.076 | < 0.001 | 0.195 | 122,598 |
| Imperial Cormorant | 1.1 ± 0.2 | 5.7 ± 0.2 | 0.190 | 0.044 | < 0.001 | 0.062 | 11,068 |
| Red-tailed Tropicbird | 1.8 ± 0.3 | 4.0 ± 0.3 | 0.527 | -0.341 | < 0.001 | 0.151 | 174,190 |
| Black-legged Kittiwake | 2.0 ± 0.4 | 4.0 ± 0.2 | 0.998 | -1.915 | < 0.001 | 0.383 | 21,767 |
| Great Frigatebird | 1.7 ± 0.3 | 2.6 ± 0.2 | 0.757 | -0.213 | < 0.001 | 0.256 | 2,805 |
| Streaked Shearwater | 1.4 ± 0.1 | 4.1 ± 0.3 | 0.018 | 1.315 | < 0.001 | 0.001 | 18,036 |
| Northern Fulmar | 1.3 ± 0.1 | 4.7 ± 0.3 | -0.003 | 1.354 | 0.437 | 0.000 | 8,505 |
| Grey-headed Albatross | 1.4 ± 0.1 | 3.1 ± 0.2 | 0.016 | 1.325 | 0.500 | -0.001 | 590 |
| Wandering Albatross | 1.1 ± 0.1 | 2.8 ± 0.2 | 0.043 | 0.952 | 0.207 | 0.001 | 533 |
| Northern Gannet | 2.5 ± 0.6 | 3.9 ± 0.3 | 0.489 | 0.632 | < 0.001 | 0.051 | 15,410 |

\*wind tunnel studies

We then examined the coefficient of variation (c.v.) in wingbeat frequency, to assess whether this varied with bird mass or morphology. These coefficients were calculated by pooling data from all individuals of the same species to cover the various flight conditions (e.g., wind speeds) experienced across tracks. None of the correlations were significant, but there was some indication that the variation in wingbeat frequency was negatively correlated with the residual wing loading (Figure 4) (Pearson’s correlation: ρ = -0.445, R^2^ = 0.131, p-value = 0.111).

**Figure 4.**
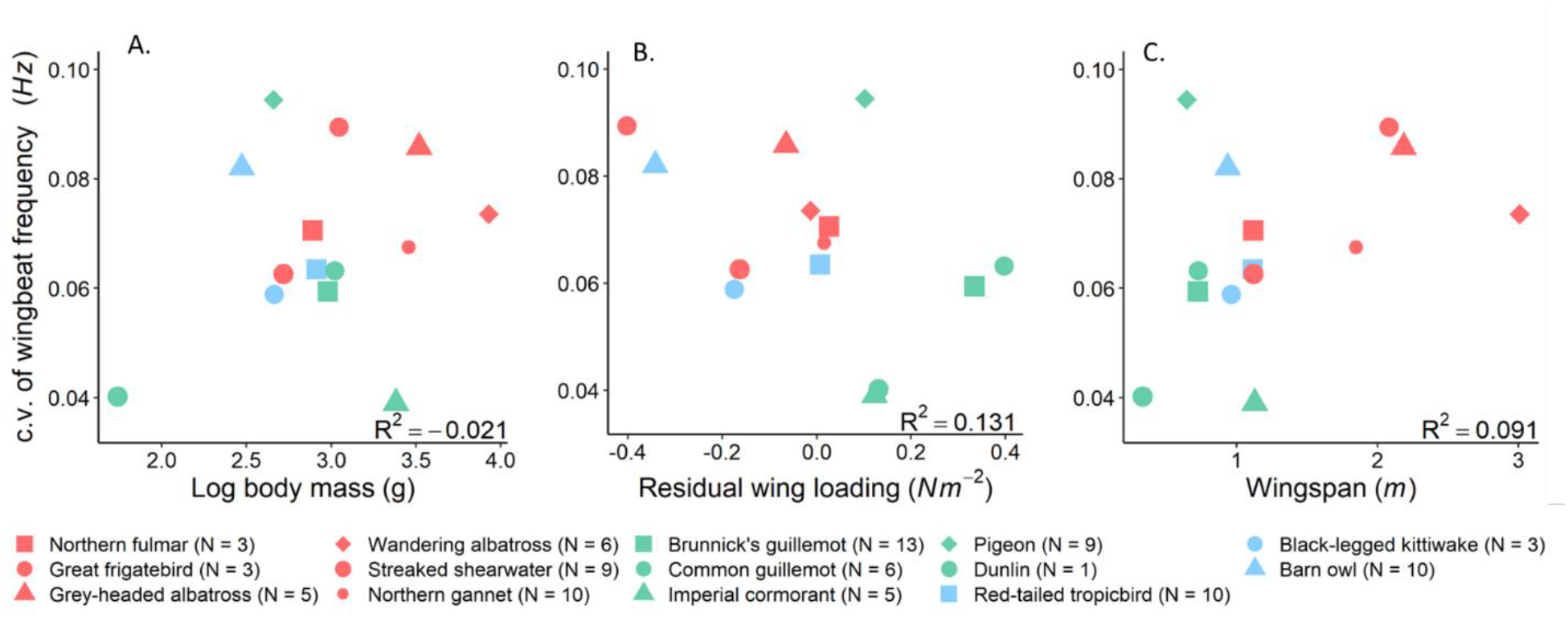
Variation in wingbeat frequency as a function of morphological parameters for 14 species: A. body mass, B. residual wing loading (where positive values indicate species with higher wing loading than expected for a given mass), and C. wingspan. Birds with similar flights style are marked with same colour: Red represents specialist soaring fliers, green represents typically flapping fliers and the blue indicates birds that use mix of flapping and soaring.

### (iii) Do birds adjust different kinematic parameters to vary speed and climb rate?

Climb rate had a positive effect on the relationship between wingbeat frequency and amplitude in tropicbirds, demonstrating that birds increased their wingbeat amplitude to a greater extent in climbing flight (Table 3). The same effect was seen in barn owls, although the R^2^ was low. We were not able to make any meaningful conclusion concerning pigeons flying in the wild as the fixed effects in the model explained only 1% of the variance in the response variable (R^2^_m_ = 0.01, see Table 3).

**Table 3.** The models of amplitude as a function of wingbeat frequency (WBF) and the interaction between wingbeat frequency and climb rate (V_z_) for red-tailed tropicbirds (n = 10), pigeons (n = 9) and barn owls (n = 10), using individual as a random factor.

|  | Estimate | Std. Error | t-value | p |
| --- | --- | --- | --- | --- |
| **A. Tropicbirds ($R^2_m = 0.50$, $R^2_c = 0.66$)** |  |  |  |  |
| (Intercept) | -2.275 | 0.056 | -40.622 | < 0.001 |
| WBF | 1.014 | 0.011 | 91.817 | < 0.001 |
| WBF: $V_z$ | 0.018 | 0.001 | 13.301 | < 0.001 |
| **B. Pigeons ($R^2_m = 0.01$, $R^2_c = 0.42$)** |  |  |  |  |
| (Intercept) | 3.882 | 0.132 | 29.524 | < 0.001 |
| WBF | -0.053 | 0.018 | -3.013 | 0.003 |
| WBF: $V_z$ | -0.008 | 0.003 | -3.256 | 0.001 |
| **C. Barn owls ($R^2_m = 0.28$, $R^2_c = 0.65$)** |  |  |  |  |
| (Intercept) | -0.615 | 0.0799 | -7.7 | < 0.001 |
| WBF | 0.677 | 0.0045 | 150.3 | < 0.001 |
| WBF: $V_z$ | 0.048 | 0.0008 | 59.3 | < 0.001 |

Airspeed did not affect the relationship between wingbeat frequency and amplitude in tropicbirds (p = 0.164), barn owls (p = 0.546), or in pigeons, where the model explained only 3% of the variability in the response variable (R^2^_m_ = 0.03, see Table SI 2). In contrast, there was a clear increase in heave amplitude with airspeed for a pigeon flying in the wind tunnel (Figure 5).

**Figure 5.**
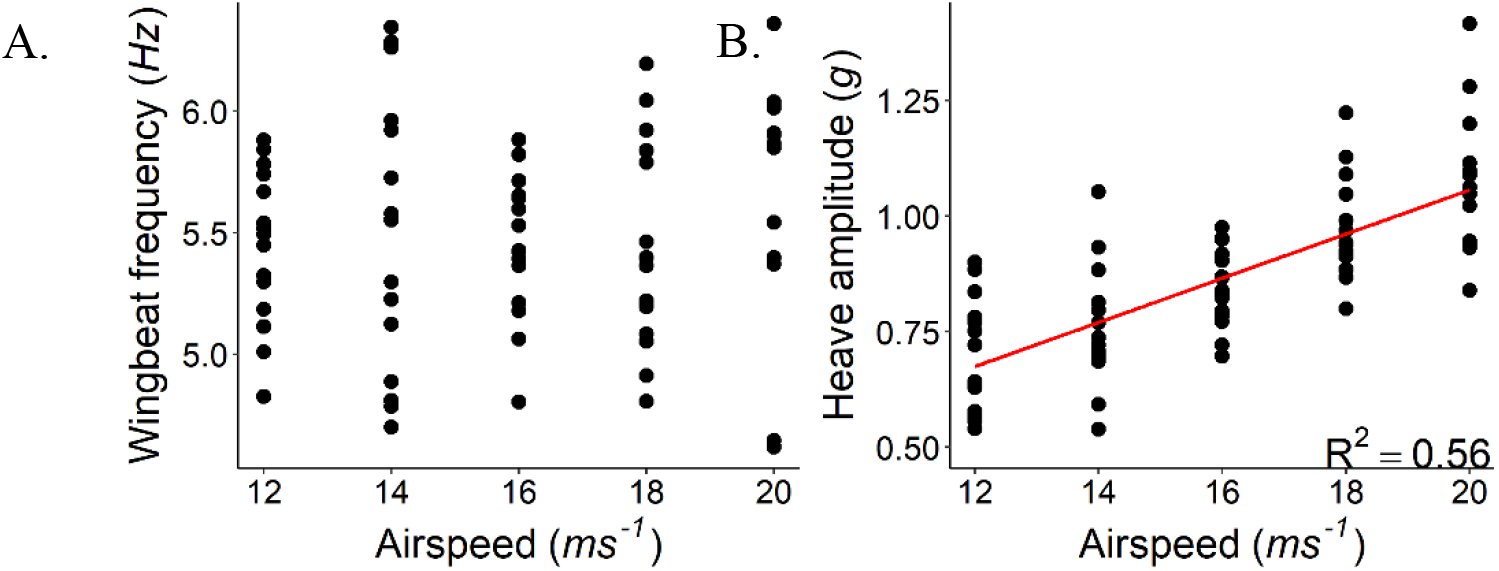
A. Wingbeat frequency and B. signal amplitude for a pigeon flying in a wind tunnel at a range of airspeeds. Each data point is an average 5 consecutive wingbeats. Periods of consistent flight were selected for analysis.

We found 22 studies where the relationship between wingbeat frequency, wingbeat amplitude and either mechanical power, speed or climb rate was quantified (Table 4a). Of these, ten were performed with Passeriformes. Kinematic analyses were mostly conducted using high speed cameras to quantify wingbeat frequency and amplitude for birds either flying in wind tunnels or flight chambers.

**Table 4a.** Summary of studies assessing the relationship between wingbeat frequency, amplitude, and mechanical power output.

| Species | Method | Flight mode | Speed (m s<sup>-1</sup>) | Remarks | Source |
| --- | --- | --- | --- | --- | --- |
| **Pigeon**<br>*Columba livia* | Field data – GPS and accelerometer measurements | Level, ascending and descending flight, all while circling | 10-18 | <u>As speed increased</u><br>WBF – varied approx. U shaped<br>WBA – increased<br><u>At constant speed, as power increases</u><br>WBF – increased<br>WBA – decreased<br><u>Ascending flight</u><br>WBF – increased<br>WBA – increased<br><u>Accelerating flight</u><br>WBF – increased<br>WBA – increased | (Usherwood et al., 2011) |
| **Pigeon**<br>*Columba livia* | Platform – muscle force measurements and kinematic analysis with high-speed cameras | Ascending, level and descending | 1.4-3.9 | <u>In different flight modes</u><br>WBF – did not vary significantly<br>WBA – decreased during take-off and prior to landing | (Tobalske & Biewener, 2008) |
| **Common starling**<br>*Sturnus vulgaris* | Wind tunnel – respirometry masks and kinematics analysis with high-speed cameras | Level flight | 6-14 | <u>As speed increased</u><br>WBF – increased (less significant)<br>WBA – increased (less significant)<br>Power – increased | (Ward et al., 2001) |
| **Eurasian tree sparrow**<br>*Passer montanus* | Experiments in flight chamber – kinematics analysis with high-speed cameras | Vertical flight | - | <u>As maximum load-lifted</u><br>WBF – no significant variation<br>WBF – no significant variation | (Wang et al., 2019) |
| **Barn swallow**<br>*Hirundo rustica* | Wind tunnel – energetic costs measured by DLW, and kinematics analysis is by video recordings | Level flight | 8-11.5 | <u>As speed increases</u><br>WBF – varied as U shaped<br><u>As mass increased</u><br>WBF – increased<br>Power - increased | Schmidt-Wellenburg et al., 2007 |
| **Blue tit**<br>*Cyanistes caeruleus* | Flight inside a custom-built box – kinematics analysis with high-speed cameras | Take-off | 3.4 | <u>As wing loading increased</u><br>WBF – decreased<br>WBA – did not vary<br>Power – decreased<br>AR – increased | (McFarlane et al., 2016) |
| **Thrush nightingale**<br>*Luscinia luscinia* | Wind tunnel – PIV and kinematics analysis with high-speed cameras | Level flight | 5-10 | <u>As speed increased</u><br>WBF – no significant variation<br>WBA – no significant variation | (Rosén et al., 2004) |
| **Thrush nightingale**<br>*Luscinia luscinia* | Wind tunnel – Wingbeat frequency measured using a shutter stroboscope and video recording | Level flight | 5-16 | <u>As mass increased</u><br>WBF – increased<br><u>As speed increased</u><br>WBF – varied in U shape (less significantly) | (Pennycuik et al., 1996) |
| **Zebra finch**<br>*Taeniopygia guttata* | Wind tunnel – kinematics analysis with high-speed cameras | Intermittent flap-bounding flight | 0-14 | <u>As speed increased</u><br>WBF – increased (less significant)<br>WBA – decreased (significantly) | (Tobalske et al., 1999) |
| **Zebra finch**<br>*Taeniopygia guttata* | Surgical procedures to measure flight muscle activity | ? | - | <u>As power increased</u><br>WBF – no significant effect<br>WBA – increased effectively | (Bahlman et al., 2020) |
| **Zebra finch**<br>*Taeniopygia guttata* | Wind tunnel – muscle *in vivo* pectoralis fascicle strain measurements, and kinematics by high-speed video recordings | Level flight | 0-14 | <u>As speed increased</u><br>WBF – varied approx. U shaped<br>WBA – increased only at hovering | (Ellerby & Askew, 2007) |
| **Budgerigar**<br>*Melopsittacus undulates* | Wind tunnel – muscle *in vivo* pectoralis fascicle strain measurements, and kinematics by high-speed video recordings | Level flight | 4-16 | <u>As speed increased</u><br>WBF – varied approx. U shaped<br>WBA – did not vary significantly | (Ellerby & Askew, 2007) |
| **Cockatiel**<br>*Nymphicus hollandicus* | Wind tunnel – *in vivo* pectoralis muscle length change measurements | Level flight | 0-16 | <u>As speed increased</u><br>Power – increased (approx. U shaped)<br>WBF – reduced (highest at the lower range) | (Morris & Askew, 2010) |
| **Cockatiel**<br>*Nymphicus hollandicus* | Wind tunnel – *in vivo* surgical procedures and kinematics analysis with high-speed cameras | Level flight | 0-14 | <u>As speed increased</u><br>WBF – reduced at lower speed and increased at higher speed (approx. U shaped)<br>Power – varied (approx. U shaped) | (Hedrick et al., 2003; Tobalske et al., 2003) |
| **Eurasian teal**<br>*Anas crecca* | Wind tunnel – Wingbeat frequency measured using a shutter stroboscope and video recording | Level flight | 5-16 | <u>As mass increased</u><br>WBF – increased<br><u>As speed increased</u><br>WBF – varied in U shape (less significantly) | (Pennycuik et al., 1996) |
| **Black-legged kittiwake**<br>*Rissa tridactyla* | Wild study – kinematics and airspeed data of commuting flights measured using GPS and accelerometer devices | Flap-glide flight (predominantly flapping) | 2-16 | <u>As speed increased</u><br>WBF – no significant relationship<br>WBA – increased significantly (as proxy by body moving amplitude) | (Collins et al., 2020) |
| **Harris's hawk**<br>*Parabuteo unicinctus* | Outdoor flight – accelerometry data and kinematic analysis using video recordings | Climbing flight | - | <u>As climb power increased</u><br>WBF – increased linearly with lesser variation<br>WBA – increased linearly with higher variation (as proxy by body moving amplitude) | (Van Walsum et al., 2020) |
| **Common swift**<br>*Apus apus* | Wind tunnel – PIV and kinematics analysis with high-speed cameras | Level flight | 8-9.2 | <u>As speed increased</u><br>WBF – decreased<br>WBA – increased | (Henningsson et al., 2008) |
| **Ruby-throated hummingbird**<br>*Archilochus colubris* | Flight experiments in an airtight cube – varying air density treated with heliox | Hovering | - | <u>As power increased</u><br>WBF – increased (less significant)<br>WBA – increased (significantly) | (Chai & Dudley, 1995) |
|  |  |  |  | <u>As air density decreased</u><br>Power – increased |  |
| **Ruby-throated hummingbird**<br>*Archilochus colubris* | Flight experiments in an airtight cube – varying air density treated with helium | Hovering | - | <u>As power increased</u><br>WBF – did not vary<br>WBA – increased (significantly) | (Chai & Dudley, 1996) |
|  |  |  |  | <u>As air density decreased</u><br>Power – increased |  |
| **Ruby-throated hummingbird**<br>*Archilochus colubris* | Cubic testing arena - surgical procedures to measure flight muscle activity and kinematics analysis with high-speed cameras | Hovering | - | <u>As load-lifted increased</u><br>WBF – did not vary<br>WBA – increased (significantly) | (Mahalingam & Welch, 2013) |
|  |  |  |  | <u>As air density decreased</u><br>WBF – did not vary<br>WBA – increased (significantly) |  |
| **Rufous hummingbird**<br>*Selasphorus rufus* | Wind tunnel – kinematics analysis with high-speed cameras | Hovering and level flight | 0-12 | <u>As speed increased</u><br>WBF – did not vary<br>WBA – increased (approx. U shaped) | (Tobalske et al., 2007) |

**Table 4b.** Summary of studies assessing the relationship between wingbeat frequency, amplitude, and metabolic power.

| Species | Method | Flight mode | Speed (m s<sup>-1</sup>) | Remarks | Source |
| --- | --- | --- | --- | --- | --- |
| **Common starling**<br>*Sturnus vulgaris* | Wind tunnel – measurements of oxygen consumption and carbon dioxide production, and kinematics analysis recorded on magnetic tape | Burst flapping and gliding | 6-18 | <u>As power stayed almost constant</u><br>WBF – constant<br>WBA – varied approx. U shaped | (Torre-Bueno & Larochelle, 1978) |
| **Black-billed magpie**<br>*Pica hudsonia* | Wind tunnel – pectoralis muscle force based on bone-strain recordings and muscle fibre length | Hovering and level flight | 0-14 | <u>As power varied L shaped</u><br>WBF – varied U shaped | (Dial et al., 1997) |
| **Cockatiel**<br>*Nymphicus hollandicus* | Wind tunnel – measurements of oxygen consumption and carbon dioxide production | Level flight | 6-14 | <u>As speed varied as U shaped</u><br>WBF – varied approx. U shaped<br>WBA – varied approx. U shaped | (Morris et al., 2010) |
| **Cockatiels**<br>*Nymphicus hollandicus* | Wind tunnel – measurement of oxygen consumption using masks | Level flight | 5-15 | <u>As power varied as U shaped</u><br>WBF – decreased significantly | (Bundle et al., 2007) |
| **Budgerigars**<br>*Melopsittacus undulatus* | Wind tunnel – measurement of oxygen consumption using masks | Level flight | 5-15 | <u>As power varied as U shaped</u><br>WBF- did not vary significantly | (Bundle et al., 2007) |
| **Budgerigars**<br>*Melopsittacus undulatus* | Wind tunnel – measurements of oxygen consumption and carbon dioxide production | Ascending, level and descending flight | 5-13 | <u>As power varied as U shaped</u><br>WBF - constant | (TUCKER, 1968) |
| **Bar-headed goose**<br>*Anser indicus* | Migratory flight – measurements using data loggers | Ascending, level and descending flight | - | Power increased as WBF<sup>6.96</sup><br>WBA increased with power | (Bishop et al., 2015) |

Wingbeat frequency had a U-shaped relationship with speed (to a variable degree) in the following species: pigeon, barn swallow, thrush nightingale, zebra finch, budgerigar and Eurasian teal. However, two further studies with thrush nightingale and cockatiel found no/ different relationships between wingbeat frequency and speed, and another three studies found no relationship between wingbeat frequency and speed in black-legged kittiwake, common swift and a rufous hummingbird (Table 4a). One study found a U-shaped relationship between wingbeat amplitude and speed, and three others found a notable positive relationship (in pigeons, kittiwake and common swift) (Table 4a). The two studies on pigeons found that wingbeat amplitude varied in ascending and descending flights. Information on wingbeat amplitude was available for the budgerigars and it did not vary significantly with speed. Four studies on hummingbirds showed that wingbeat amplitude increased with power during hovering.

Results from a further seven studies showed that the relationships between metabolic power and wingbeat frequency and amplitude were similarly variable (Table 4b). While wingbeat frequency was positively correlated with the metabolic power in four studies, either it did not vary significantly or stayed constant in two studies, and it declined in one study. Furthermore, cockatiels in two different studies exhibited a discrepancy between the wingbeat frequency and the power variation for the same speed range and flight mode: While the power had a U-shaped relationship with speed, wingbeat frequency varied in same fashion in one case but was negatively related to speed in another. Out of three studies reported, wingbeat amplitude was closely correlated with power in two.

## Discussion

The total power output of a bird in flapping flight varies between level, accelerating, ascending/descending, manoeuvring and load carrying flight, as well as with flight speed. Birds are expected to modulate the power output predominantly through wingbeat frequency and/ or wingbeat amplitude changes, as first principles state that power output is directly proportional to the cube of the product of wingbeat frequency and amplitude. Metrics from onboard accelerometers should be able to provide insight into the relative importance of both these parameters. Our data from wind tunnel flights confirm this, by showing that the amplitude of the dorsoventral body acceleration (heave) and the wingbeat amplitude are positively related within a wingbeat cycle. While the R^2^ values varied substantially between the two pigeons (0.08 versus 0.24, see Figure 3), this is unlikely to reflect differences in kinematics, which should be consistent across individuals for the same flight style, instead, the variance in these relationships is likely to have been affected by the flight consistency, and possibly the stability of the magnet attachment. More broadly, the ability to resolve relative changes in wingbeat amplitude from the acceleration signal may show some variation with flight style, for instance, peaks associated with wingbeats can be harder to resolve against a baseline that varies due to centripetal acceleration, as occurs throughout the dynamic soaring cycle. This may help explain the lack of a correlation between wingbeat frequency and acceleration amplitude in three of the four species that used dynamic soaring in this study, although this could also reflect a genuine absence of a relationship in this group.

The question that follows is, to what extent do birds modify wingbeat frequency and/ or amplitude to modulate power output? We found that wingbeat frequency and amplitude were correlated for pigeons, and wingbeat amplitude increased with increasing flight speed (similar to another pigeon study, Usherwood et al., 2011). It was therefore surprising that we found no relationship between wingbeat frequency, amplitude and airspeed in pigeons during homing flights. The discrepancy between our wind tunnel and “wild” flights may be related to the extremely variable nature of pigeon homing flights when flying solo (Garde et al., 2021). Indeed, the substantial (and costly) variation in speed and rate of change in altitude has been proposed to serve as a predator avoidance strategy, which birds such as pigeons may adopt when flocking is not possible (Garde et al., 2021). This is relevant in the current context as it could mask a relationship between wingbeat frequency, amplitude and airspeed in homing flights. This therefore highlights that birds experience very different biological and physical environments when flying in the laboratory and in the wild, which can in turn affect their kinematics. There are also likely to be errors in the estimation of airspeed, as wind conditions were recorded near the release site and while this was within 5.7 km of the loft, the wind field will be affected by the local topography as well as flight altitude. These errors will be larger for the tropicbird study, where GPS locations were recorded once a minute and wind speeds were measured up to tens of kilometres away from the bird locations, which likely contributes to the lack of a correlation between kinematic parameters and airspeed in this species. Nonetheless, the positive relationship between kinematic parameters and climb rates for tropicbirds shows that relationships can be resolved using high frequency data from birds flying in the wild as, unlike wind, pressure was recorded with sub-second resolution.

Our finding that wingbeat frequency and amplitude were positively correlated in 11 of the 14 species that we investigated suggests that both parameters tend to be involved in power modulation across a range of morphologies and body masses. However, the low R^2^ values indicate that they are unlikely to covary in a straightforward manner, as also indicated by the variable relationship between wingbeat frequency and amplitude in other studies: five studies reported a positive correlation, five reported a negative relationship and three reported no correlation (Table 2). Nonetheless, our review of the literature did suggest that birds tend to increase their wingbeat amplitude more in the most energetically demanding forms of flight (Table 4a), consistent with our finding that tropicbirds increased their wingbeat amplitude to a greater extent than frequency when climbing. For instance, while Usherwood *et al*. (2011) found that wingbeat frequency increased during all flight modes for pigeons flying in a flock, the wingbeat amplitude increased with induced power, climb rate, and accelerating flight. Parallels can be found in studies by Tobalske and Biewener (2008), where pigeons varied their wingbeat amplitude, but not frequency, during take-off and landing. Zebra finches (*Taeniopygia guttata*) were also found to modulate wingbeat amplitude rather than wingbeat frequency for high power events (Bahlman et al., 2020, Ellerby & Askew, 2007), but not in level flight (Ellerby & Askew, 2007). Other studies have shown that wingbeat amplitude increased to meet the power demand associated with load carrying in hovering/vertical flight, whereas the wingbeat frequency remained near constant (Mahalingam & Welch, 2013). Similarly, hummingbirds increased their wingbeat amplitude when flying in low density air, both in the laboratory (Chai & Dudley, 1995; Chai & Dudley, 1996; Mahalingam & Welch, 2013) and in the field along natural elevational gradients (Altshuler & Dudley, 2006; Altshuler & Dudley, 2003), with wingbeat amplitudes up to 180° at flight failure densities.

Nonetheless, flight mode alone does not explain which kinematic parameter birds select to modulate their flight power, as while we found 10 studies where wingbeat frequency increased with airspeed in non-hovering flight, there were negative relationships between frequency and airspeed in two studies, and no relationship in ten studies (Table 4) including our data. The variation across studies is striking and extends beyond comparisons between laboratory and field settings. In fact differing relationships were found in two species (cockatiels and thrush nightingales) in experimental studies, which may indicate the role of factors such as turbulence levels in wind tunnels, or the difficulties of training birds to maintain steady level flight, both of which could have a notable impact on the variability of kinematic parameters over fine scales.

We found limited support for the hypothesis that morphology influences variation in kinematic parameters, although birds with high residual wing loading, such as auks, did appear to have relatively low variation in wingbeat frequency, consistent with their relatively low available power. It would be interesting to see whether this non-significant negative correlation persists with data from a greater number of species.

This study has focused on variation in wingbeat frequency and amplitude. However, birds can also vary the aerodynamic forces through changes in the other wingbeat kinematic parameters and wing flexing and it is unclear whether and how they could all be captured by body-mounted accelerometers. Other kinematics parameters that have a significant role in power output include the upstroke-to-downstroke ratio, stroke-plane angle, span ratio, twist, and angle of attack. In experiments with a house martin (*Delichon urbicum*) and a thrush nightingale (*Luscinia luscinia*), the upstroke-to-downstroke ratio and span ratio varied with increasing flight speed, whereas the wingbeat frequency and amplitude did not (Rosén et al., 2004; Rosén et al., 2007). Similarly, Ward et al. (2001) showed that for a common starling (*Sturnus vulgaris*), the wingbeat frequency and amplitude were the least important parameters associated with an increase in power, compared to variations in the stroke-plane angle and downstroke ratio. Finally, several species vary the body angle and stroke-plane angle to support weight at low speeds and augment thrust at higher speeds, while frequency and amplitude varied to a lesser degree in these scenarios (Tobalske & Dial, 1996). The situation is potentially even more complex in intermittent flap-bounding flight, and indeed, cycle time spent flapping, flapping-and-bounding duration, and the number of flaps were more important than wingbeat frequency and amplitude for a zebra finch increasing its flight speed (Tobalske et al., 1999).

Overall, in terms of the implications for acceleration metrics to act proxies for flight power, it is clear that body mounted accelerometers can provide information on wingbeat amplitude as well as frequency, both of which show substantial variation when considered across free-ranging flights in multiple species. Acceleration metrics that incorporate variation due to wingbeat frequency and amplitude, such as DBA and body power (Spivey & Bishop, 2013; Wilson et al., 2006) should therefore be more robust proxies for power use than wingbeat frequency alone. In support of this, DBA has been shown to be a better predictor of overall energy expenditure (estimated with doubly labelled water) than flight time or wingbeat frequency in auks (Elliott et al., 2013; Elliott et al., 2014). Nonetheless, wingbeat frequency and amplitude are only partial determinants of the wingbeat kinematics associated with power, and other factors play a substantial role in power production for certain flight types (Berg & Biewener, 2008). Some of these e.g., the downstroke ratio, may be estimated from onboard accelerometers (Taylor et al., 2019), although the magnetometer is a valuable addition in this regard, highlighting when the downstroke begins and ends (e.g., Figure 2). Beyond this, what is clear is that while relationships between DBA and energy expenditure are linear for terrestrial and aquatic forms of locomotion (a relationship that holds across tens of species and over different timeframes, (Halsey et al., 2009; Halsey et al., 2011; Wilson et al., 2020)), it is unlikely to be the case for all types of flight, not least because of the varying contribution of wingbeat frequency and amplitude to power. Experiments with independent estimates of power output will provide further insight into the performance of acceleration-based proxies and the extent to which single metrics applied across species and contexts.

## Acknowledgements

This work was supported by a European Research Council starter grant 715874 to ELCS, under the European Union’s Horizon 2020 research and innovation program. Rory Wilson supplied the tags that were used for data collection in the studies on Imperial cormorants, wandering albatrosses, grey-headed albatrosses and streaked shearwaters. Fieldwork on northern fulmars was supported by the BlueFish project, funded by the European Regional Development fund through the Ireland Wales Co-operation Programme (2014−2020) and an extended field team. Fieldwork on northern gannets was supported by the FishKOSM project funded by the Department of Agriculture Food and the Marine (15/S/744). The wind tunnel experiments at Lund University were supported by the Swedish Research Council (2016-03625) and A.H. Linus Hedh helped training the dunlin for wind tunnel flight. Data collection in the Max Planck wind tunnel was supported by a Max Planck Sabbatical Fellowship (to ELCS). Fieldwork on black-legged kittiwakes was assisted by the Middleton Island Field Crew of 2019. Funding for Brunnich’s guillemot and black-legged kittiwake work was partially from the Natural Sciences and Engineering Research Council of Canada (to KE). Field work on the barn owls was funded by Swiss National Science Foundation (310030, 200321 to AR). We thank the National Parks and Conservation Service, Government of Mauritius, for permission to conduct the field work on the red-tailed tropicbirds and the Round Island Wardens for their support in the field. Research in Japan was funded by the Grants-in-Aid for Scientific Research from the Japan Society of the Promotion of Science (16K21735, 16H06541, 21H05294 to KY). We also thank Gil Bohrer for essential help in setting up the sonic anemometer for in-flight measurements of air movement.

## Supplementary Information

**Table SI 1.** Ethical approvals for the data used for analysis in this manuscript.

| Species | Location | Ethical approval reference |
| --- | --- | --- |
| **Brünnich's guillemot**<br>*Uria lomvia* | Coats Island, Nunavut, Canada | Approved under Environment Canada Animal Care permits 1000-AG01a, 11-AG02, EC-PN-12-0, Migratory Bird Research Permits NUN-MBS-09-01, NUN-SCI-11-07 and NUN-SCI-12-01 and Nunavut and Nunavik Wildlife Research permits (WL-2010-038; WL-2011-019, WL-2012-04-06). |
| **Common guillemot**<br>*Uria aalge* | Puffin Island, UK | Swansea University AWERB, permit SU-Ethics-Staff-050619/150 (reference number: STAFF_BIOL_25956_280519125243_1). |
| **Northern fulmar**<br>*Fulmarus glacialis* | Saltee Islands, Ireland | UCC Animal Ethics Committee approval. Work conducted under licences from the British Trust for Ornithology (BTO) and Irish National Parks and Wildlife Service (NPWS). |
| **Pigeon**<br>*Columba livia* | Radolfzell, Germany (free flight) | Swansea University AWERB, issue number IP-1718-23 and Regierungspräsidium Freiburg, Baden-Württemberg, Germany, permit number G-17/92 |
| **Pigeon**<br>*Columba livia* | Seewiesen, Germany (wind tunnel) | Approved by the government of Upper Bavaria, "Sachgebiet 54 – Verbraucherschutz, Veterinärwesen, 80538 München" with the record number: Gz.: 55.2-1-54-2532-86-2015. |
| **Red-tailed tropicbird**<br>*Phaethon rubricauda* | Round Island, Mauritius | Swansea University AWERB, permit 040118/39. |
| **Great frigatebird**<br>*Fregata minor* | Europa Island | Approved by the Préfet des Terres Australes et Antarctiques Françaises and Comité National de la Protection de la Nature. |
| **Black-legged kittiwake**<br>*Rissa tridactyla* | Middleton Island, Alaska, USA | Swansea University AWERB, permit 110619/1590 (IP-1819-18). |
| **Imperial cormorant**<br>*Leucocarbo atriceps* | Punta Leon, Argentina | Approved by the Dirección de Flora y Fauna Silvestre de la Provincia de Chubut and the Organismo Provincial de Turismo de Chubut. |
| **Western barn owl**<br>*Tyto alba* | Switzerland | Legal authorisations VD 2844, VD 3213 and VD 3571. |
| **Grey-headed albatross**<br>*Thalassarche chrysostoma* | Marion Island, South Africa | Approved by the University of Cape Town's Science Faculty Animal Ethics Committee (2017/V10REV/PRyan). |
| **Wandering albatross**<br>*Diomedea exulans* | Marion Island, South Africa | Approved by the University of Cape Town's Science Faculty Animal Ethics Committee (2017/V10REV/PRyan). |
| **Streaked shearwater**<br>*Calonectris leucomelas* | Awashima Island, Japan | Approved by the Animal Experimental Committee of Nagoya University and the Ministry of the Environment Government of Japan. |
| **Dunlin**<br>*Calidris alpina* | Sweden | Swansea University AWERB, permit number 030718/66. |
| **Northern gannet**<br>*Morus bassanus* | Saltee Islands, Ireland | UCC Animal Ethics Committee approval. Work conducted under licences from the British Trust for Ornithology (BTO) and Irish National Parks and Wildlife Service (NPWS: C87/2017, C63/2018). |

**Table SI 2.** Output of the model of amplitude as a function of wingbeat frequency and the interaction between wingbeat frequency and airspeed in level flapping flight for red-tailed tropicbirds (n = 10), pigeons (n = 9) and western barn owls (n = 10), using individual as a random factor.

|  | Estimate | Std. Error | t-value | p |
| --- | --- | --- | --- | --- |
| **A. Tropicbirds ($R^2_m = 0.11$, $R^2_c = 0.14$)** |  |  |  |  |
| (Intercept) | 1.137 | 0.377 | 3.016 | 0.004 |
| WBF | 0.273 | 0.095 | 2.876 | 0.006 |
| WBF: Airspeed | -0.003 | 0.002 | -1.41 | 0.164 |
| **B. Pigeons ($R^2_m = 0.03$, $R^2_c = 0.42$)** |  |  |  |  |
| (Intercept) | 3.714 | 0.229 | 16.198 | < 0.001 |
| WBF | -0.143 | 0.053 | -2.688 | 0.008 |
| WBF: Airspeed | 0.006 | 0.002 | 3.95 | < 0.001 |
| **C. Barn owls ($R^2_m = 0.25$, $R^2_c = 0.62$)** |  |  |  |  |
| (Intercept) | -0.533 | 0.070 | -7.6 | < 0.001 |
| WBF | 0.658 | 0.005 | 133.9 | < 0.001 |
| WBF: Airspeed | 0 | 0 | 0.6 | 0.546 |

### Effect of sampling frequency on the estimation of signal amplitude

We performed the following simulation to check whether the variation in signal amplitude could have occurred as an artefact of our sampling frequencies being similar across species, but wingbeat frequency varying. In other words, how does the decreasing number of datapoints per wingbeat cycle affect the magnitude of and variation in estimates of signal amplitude?

#### Case 1: What sampling frequency is required to minimise error in signal amplitude estimates (assessed with constant wingbeat frequency and varying sampling frequency)?

We generated a sinusoidal heaving motion with total heaving amplitude 5 g and cycle frequency with 5 Hz. The sinusoidal heaving motion was described using:

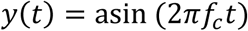

where a – is the amplitude of sinusoidal motion, *f*_*c*_– is the cycle frequency and t – is the time. If the sampling frequency is *f*_*s*_, then the wave is sampled discretely in the sampling interval of dt *= 1/f*_*s*_.

In figure 1, the mean amplitude of the motion is shown against different sampling frequencies when sampled at different product of cycle frequency (i.e., *f*_*s*_ = 1*f*_*c*_, 2*f*_*c*_, 3*f*_*c*_………n*f*_*c*_) for a period of 5 s. We can see that when sampled with one or two times the cycle frequency, the amplitude of the heaving motion is not resolved. As the sampling frequency increased, the estimated amplitude gets closer to the true amplitude (5 g). Yet, there are still some smaller deviations from the true amplitude when sampled at higher rate.

**Figure S1.**
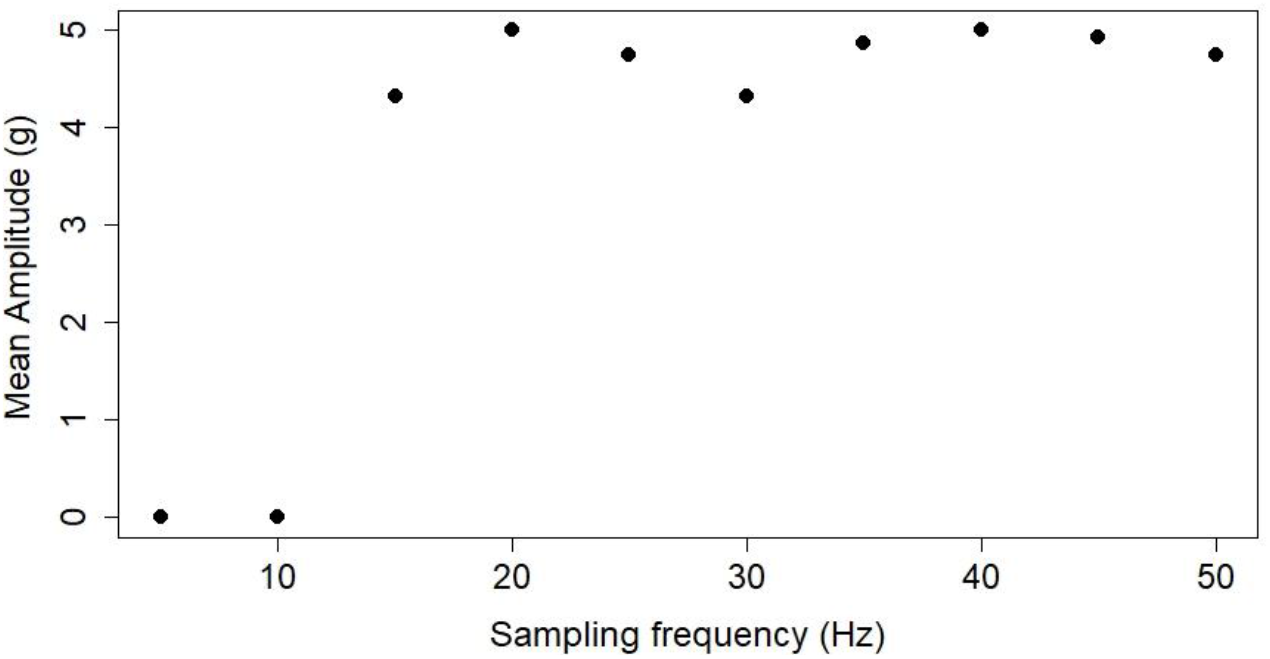
Mean estimated amplitude for a sine wave with a frequency of 5 Hz that is sampled with varying frequencies from 1f_c_to 10f_c_.

This is because a sine wave with a frequency 5 Hz has a period of 0.2 s, a positive peak at 0.05 s and a negative peak at 0.15 s. The resolved wave does not capture the exact peak if the sampled timestep occurs before/ after it. For example, when sampling with 40 Hz (8 times the cycle frequency) the time increases in 0.025 s steps as [0s:0.025s:5s] which has a discrete time point to capture the exact peak. But when sampled at 50 Hz (10 times cycle frequency) the time increases in 0.02 s increments which has a discrete time point close to the first peak (0.06 s but not 0.05 s). Thus, even if sampled at higher rate the resolved peak is underestimated depending on the cycle frequency and the sampling frequency. A generic rule in signal processing is that the waveform should be sampled at least at 10 times the cycle frequency for minimal error in peak amplitude estimation (Figure 2). This is unlikely to have caused systematic error in our study, as the wingbeats of most species were sampled with around 10 times the cycle frequency, with the only exception being the common guillemot, which was sampled with 5 times the cycle frequency.

**Figure S2.**
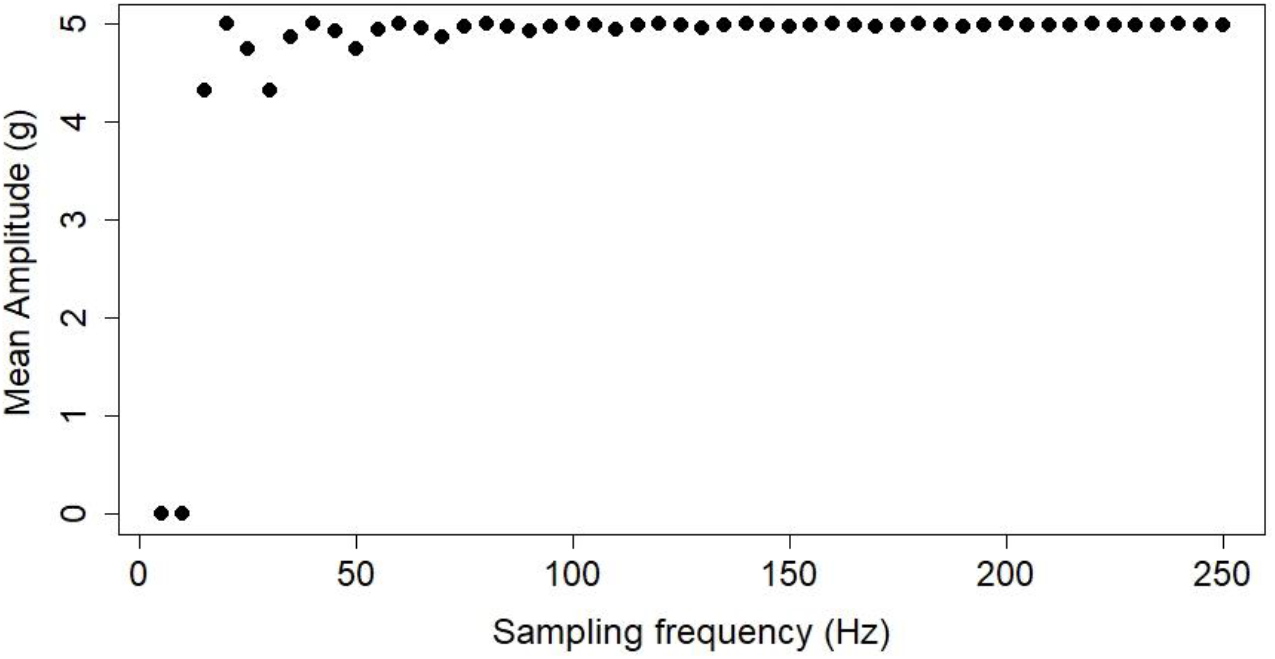
Mean estimated amplitude for a sine wave with a frequency of 5 Hz that is sampled with varying frequencies from 1f_c_to 50f_c_.

#### Case 2: How does the variation in estimated signal amplitude vary with wingbeat frequency? Assessed with a constant sampling frequency

Higher wingbeat frequencies will become increasingly under-sampled for a fixed sampling frequency (Figure 3). This should result in an increase in the variation of any mean amplitude estimate as wingbeat frequency increases. We see that for a simulated case where the signal is sampled at 40 Hz, the standard deviation increases dramatically for wingbeat frequencies of 12 Hz (Figure 4). Given that the only species with a wingbeat frequency > 10 Hz in this study was sampled at 100 Hz, we conclude that our estimates of wingbeat amplitude and associated standard deviations will not be influenced by sampling considerations.

**Figure S3.**
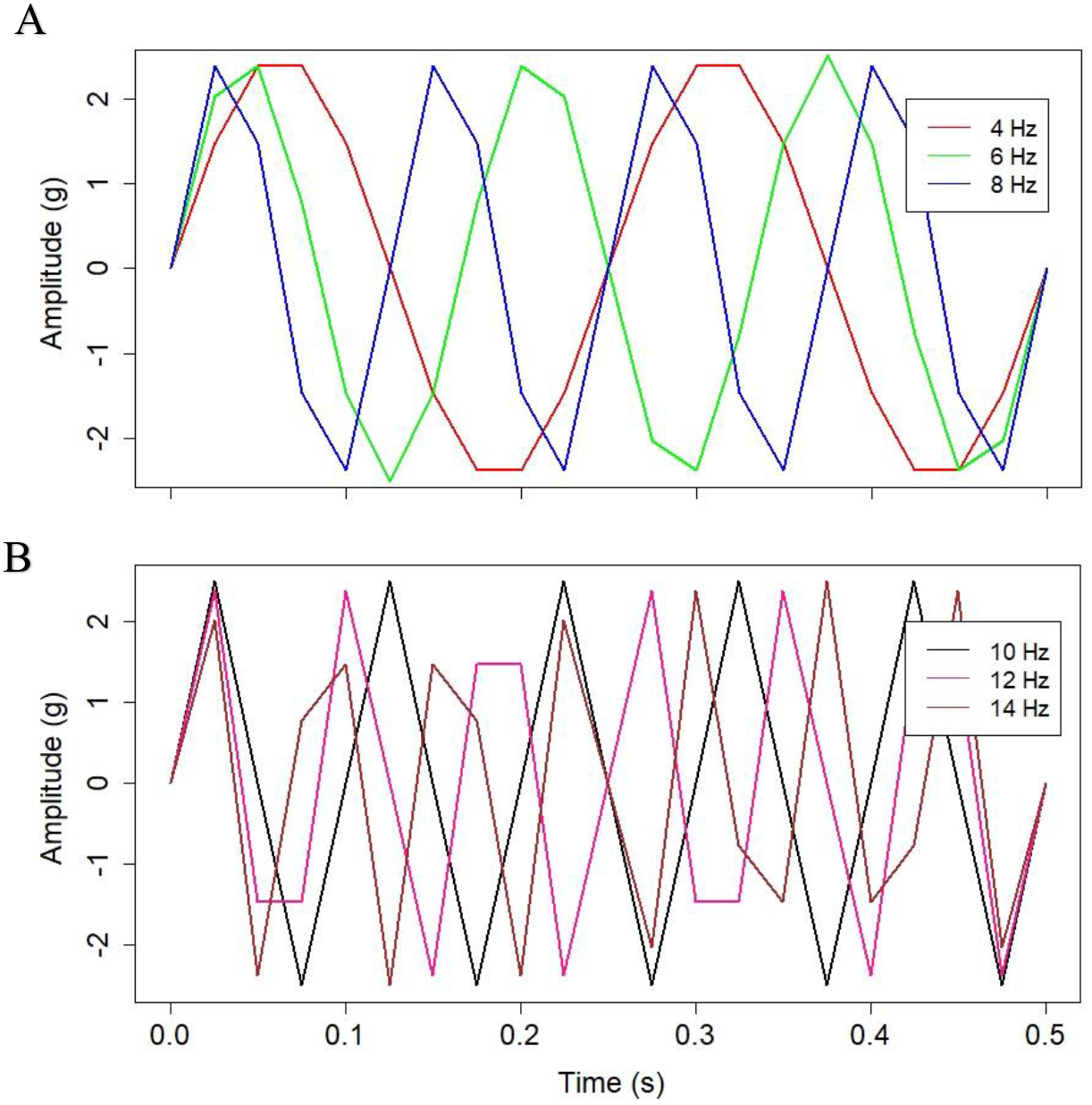
Sine waves sampled at 40 Hz with frequencies of A. 4, 6, and 8 Hz; B. 10, 12 and 14 Hz. The 10 Hz cycle shows a peculiar case where the frequency and amplitude are resolved to the true value, but the reconstructed waveform is not ideally sinusoidal because the sampling time points fall exactly on the start, peak and end times.

**Figure S4.**
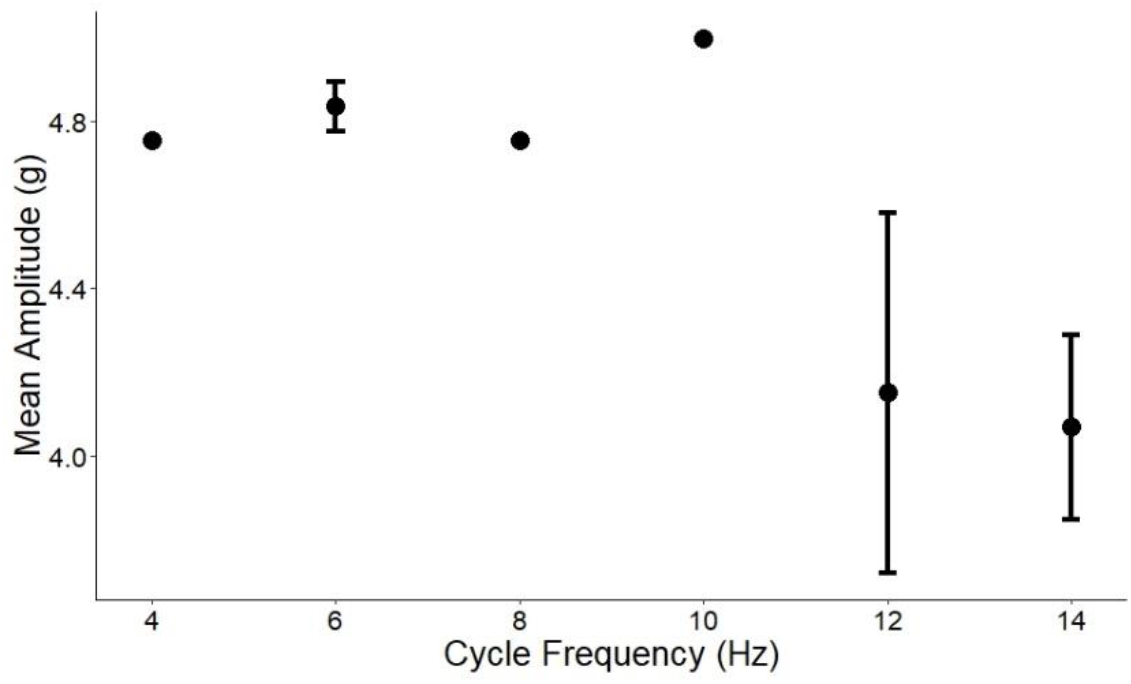
Mean amplitude with standard deviation for different cycle frequencies sampled with 40 Hz.

## References

Ainley, D. G., Porzig, E., Zajanc, D., & Spear, L. B. (2015). Seabird flight behavior and height in response to altered wind strength and direction. Marine Ornithology, 43(1), 25–36.

Altshuler, D. L., & Dudley, R. (2006). The physiology and biomechanics of avian flight at high altitude. Integrative and Comparative Biology, 46(1), 62–71. https://doi.org/10.1093/icb/icj008

Altshuler, Douglas L, & Dudley, R. (2003). Kinematics of hovering hummingbird flight along simulated and natural elevational gradients. Journal of Experimental Biology, 206(18), 3139–3147. https://doi.org/10.1242/jeb.00540

Bahlman, J. W., Baliga, V. B., & Altshuler, D. L. (2020). Flight muscle power increases with strain amplitude and decreases with cycle frequency in zebra finches (Taeniopygia guttata). Journal of Experimental Biology, 223(21), jeb225839. https://doi.org/10.1242/jeb.225839

Barton, K., & Barton, M. K. (2015). Package ‘MuMIn.’ Version, 1, 18. https://cran.r-project.org/package=MuMIn

Berg, A. M., & Biewener, A. A. (2008). Kinematics and power requirements of ascending and descending flight in the pigeon (Columba livia). Journal of Experimental Biology, 211(7), 1120–1130. https://doi.org/10.1242/jeb.010413

Bilo, D., Lauck, A., & Nachtigall, W. (1984). Measurement of linear body accelerations and calculation of the instantaneous aerodynamic lift and thrust in a pigeon flying in a wind tunnel. Biona-report, 3, 87–108.

Biro, D., Guilford, T., Dell’Omo, G., & Lipp, H. P. (2002). How the viewing of familiar landscapes prior to release allows pigeons to home faster: Evidence from GPS tracking. Journal of Experimental Biology, 205(24), 3833–3844. https://doi.org/10.1242/jeb.205.24.3833

Bishop, C. M., Spivey, R. J., Hawkes, L. A., Batbayar, N., Chua, B., Frappell, P. B., Milsom, W. K., Natsagdorj, T., Newman, S. H., Scott, G. R., Takekawa, J. Y., Wikelski, M., & Butler, P. J. (2015). The roller coaster flight strategy of bar-headed geese conserves energy during Himalayan migrations. Science, 347(6219), 250–254. https://doi.org/10.1126/science.1258732

Bundle, M. W., Hansen, K. S., & Dial, K. P. (2007). Does the metabolic rate-flight speed relationship vary among geometrically similar birds of different mass? Journal of Experimental Biology, 210(6), 1075–1083. https://doi.org/10.1242/jeb.02727

Chai, Peng, & Dudley, R. (1995). Limits to vertebrate locomotor energetics suggested by hummingbirds hovering in heliox. Nature, 377(6551), 722–725. https://doi.org/10.1038/377722a0

Chai, P., & Dudley, R. (1996). Limits to flight energetics of hummingbirds hovering in hypodense and hypoxic gas mixtures. Journal of Experimental Biology, 199(10), 2285–2295. https://doi.org/10.1242/jeb.199.10.2285

Cochran, W. W., Bowlin, M. S., & Wikelski, M. (2008). Wingbeat frequency and flap-pause ratio during natural migratory flight in thrushes. Integrative and Comparative Biology, 48(1), 134–151. https://doi.org/10.1093/icb/icn044

Collins, P. M., Green, J. A., Elliott, K. H., Shaw, P. J. A., Chivers, L., Hatch, S. A., & Halsey, L. G. (2020). Coping with the commute: behavioural responses to wind conditions in a foraging seabird. Journal of Avian Biology, 51(4), 1–11. https://doi.org/10.1111/jav.02057

Crandell, K. E., & Tobalske, B. W. (2011). Aerodynamics of tip-reversal upstroke in a revolving pigeon wing. Journal of Experimental Biology, 214(11), 1867–1873.

Delignette-Muller, M. L., Dutang, C., Pouillot, R., Denis, J.-B., & Delignette-Muller, M. M. L. (2015). Package ‘fitdistrplus.’ R Foundation for Statistical Computing, Vienna, Austria.. https://doi.org/10.18637/jss.v064.i04

Dial, K. P., Biewener, A. A., Tobalske, B. W., & Warrick, D. R. (1997). Mechanical power output of bird flight. Nature, 390(6655), 67–70. https://doi.org/10.1038/36330

Ellerby, D. J., & Askew, G. N. (2007). Modulation of pectoralis muscle function in budgerigars Melopsitaccus undulatus and zebra finches Taeniopygia guttata in response to changing flight speed. Journal of Experimental Biology, 210(21), 3789–3797. https://doi.org/10.1242/jeb.006296

Elliott, Kyle H, Le Vaillant, M., Kato, A., Speakman, J. R., & Ropert-Coudert, Y. (2013). Accelerometry predicts daily energy expenditure in a bird with high activity levels. Biology Letters, 9(1), 20120919. https://doi.org/10.1098/rsbl.2012.0919

Elliott, Kyle Hamish, Chivers, L. S., Bessey, L., Gaston, A. J., Hatch, S. A., Kato, A., Osborne, O., Ropert-Coudert, Y., Speakman, J. R., & Hare, J. F. (2014). Windscapes shape seabird instantaneous energy costs but adult behavior buffers impact on offspring. Movement Ecology, 2(1), 1–15. https://doi.org/10.1186/s40462-014-0017-2

Engel, S., Bowlin, M. S., & Hedenström, A. (2010). The role of wind-tunnel studies in integrative research on migration biology. Integrative and Comparative Biology, 50(3), 323–335. https://doi.org/10.1093/icb/icq063

Floryan, D., Van Buren, T., & Smits, A. J. (2018). Efficient cruising for swimming and flying animals is dictated by fluid drag. Proceedings of the National Academy of Sciences of the United States of America, 115(32), 8116–8118. https://doi.org/10.1073/pnas.1805941115

Garde, B., Wilson, R. P., Lempidakis, E., Börger, L., Portugal, S. J., Hedenström, A., Dell’Omo, G., Quetting, M., Wikelski, M., & Shepard, E. L. C. (2021). Fine-scale changes in speed and altitude suggest protean movements in homing pigeon flights. Royal Society Open Science, 8(5). https://doi.org/10.1098/rsos.210130

Garde, B., Wilson, R. P., Fell, A., Cole, N., Tatayah, V., Holton, M. D., Rose, K. A. R., Metcalfe, R. S., Robotka, H., Wikelski, M., Tremblay, F., Whelan, S., Elliott, K. H., & Shepard, E. L. C. (2022). Ecological inference using data from accelerometers needs careful protocols. Methods in Ecology and Evolution, 13(4), 813–825. https://doi.org/10.1111/2041-210X.13804

Halsey, L. G., Shepard, E. L. C., Quintana, F., Gomez Laich, A., Green, J. A., & Wilson, R. P. (2009). The relationship between oxygen consumption and body acceleration in a range of species. Comparative Biochemistry and Physiology - A Molecular and Integrative Physiology, 152(2), 197–202. https://doi.org/10.1016/j.cbpa.2008.09.021

Halsey, L. G., Shepard, E. L. C., & Wilson, R. P. (2011). Assessing the development and application of the accelerometry technique for estimating energy expenditure. Comparative Biochemistry and Physiology - A Molecular and Integrative Physiology, 158(3), 305–314. https://doi.org/10.1016/j.cbpa.2010.09.002

Hedenström, A., & Lindström, Å. (2017). Wind tunnel as a tool in bird migration research. Journal of Avian Biology, 48(1), 37–48. https://doi.org/10.1111/jav.01363

Hedrick, T. L., Tobalske, B. W., & Biewener, A. A. (2003). How cockatiels (Nymphicus hollandicus) modulate pectoralis power output across flight speeds. Journal of Experimental Biology, 206(8), 1363–1378. https://doi.org/10.1242/jeb.00272

Henningsson, P., Spedding, G. R., & Hedenstrom, A. (2008). Vortex wake and flight kinematics of a swift in cruising flight in a wind tunnel. Journal of Experimental Biology, 211(5), 717–730. https://doi.org/10.1242/jeb.012146

Hentze, N. T. (2012). Characteristics of over-ocean flocking by Pacific dunlins (Calidris alpina pacifica). M.Sc. Thesis, Biological Sciences Department, SFU, Canada.

Hicks, O., Burthe, S., Daunt, F., Butler, A., Bishop, C., & Green, J. A. (2017). Validating accelerometry estimates of energy expenditure across behaviours using heart rate data in a free-living seabird. Journal of Experimental Biology, 220(10), 1875–1881. https://doi.org/10.1242/jeb.152710

Kranstauber, B., Weinzierl, R., Wikelski, M., & Safi, K. (2015). Global aerial flyways allow efficient travelling. Ecology Letters, 18(12), 1338–1345. https://doi.org/10.1111/ele.12528

Lee, S. Y., Scott, G. R., & Milsom, W. K. (2008). Have wing morphology or flight kinematics evolved for extreme high altitude migration in the bar-headed goose? Comparative Biochemistry and Physiology Part C: Toxicology & Pharmacology, 148(4), 324–331. https://doi.org/10.1016/j.cbpc.2008.05.009

Mahalingam, S., & Welch Jr, K. C. (2013). Neuromuscular control of hovering wingbeat kinematics in response to distinct flight challenges in the ruby-throated hummingbird, Archilochus colubris. Journal of Experimental Biology, 216(22), 4161–4171. https://doi.org/10.1242/jeb.089383

McFarlane, L., Altringham, J. D., & Askew, G. N. (2016). Intra-specific variation in wing morphology and its impact on take-off performance in blue tits (Cyanistes caeruleus) during escape flights. Journal of Experimental Biology, 219(9), 1369–1377. https://doi.org/10.1242/jeb.126888

Morris, C. R., & Askew, G. N. (2010). The mechanical power output of the pectoralis muscle of cockatiel (Nymphicus hollandicus): The in vivo muscle length trajectory and activity patterns and their implications for power modulation. Journal of Experimental Biology, 213(16), 2770–2780. https://doi.org/10.1242/jeb.035691

Morris, C. R., Nelson, F. E., & Askew, G. N. (2010). The metabolic power requirements of flight and estimations of flight muscle efficiency in the cockatiel (Nymphicus hollandicus). Journal of Experimental Biology, 213(16), 2788–2796. https://doi.org/10.1242/jeb.035717

Norberg, U. M. (2012). Vertebrate flight: mechanics, physiology, morphology, ecology and evolution (Vol. 27). Springer Science & Business Media. https://doi.org/10.1007/978-3-642-83848-4

Orben, R. A., Paredes, R., Roby, D. D., Irons, D. B., & Shaffer, S. A. (2015). Body size affects individual winter foraging strategies of thick-billed murres in the Bering Sea. Journal of Animal Ecology, 84(6), 1589–1599. https://doi.org/10.1111/1365-2656.12410

Pennycuick, C. (1997). Actual and’optimum’flight speeds: field data reassessed. Journal of Experimental Biology, 200(17), 2355–2361. https://doi.org/10.1242/jeb.200.17.2355

Pennycuick, C. J., Alerstam, T., & Hedenström, A. (1997). A new low-turbulence wind tunnel for bird flight experiments at Lund University, Sweden. Journal of Experimental Biology, 200(10), 1441–1449. https://doi.org/10.1242/jeb.200.10.1441

Pennycuick, C. J., Hedenström, A., & Rosén, M. (2000). Horizontal flight of a swallow (Hirundo rustica) observed in a wind tunnel, with a new method for directly measuring mechanical power. Journal of Experimental Biology, 203(11), 1755–1765. https://doi.org/10.1242/jeb.203.11.1755

Pennycuick, C. J., Klaassen, M., Kvist, A., & Lindström, Å. (1996). Wingbeat frequency and the body drag anomaly: Wind-tunnel observations on a thrush nightingale (Luscinia Luscinia) and a teal (Anas crecca). Journal of Experimental Biology, 199(12), 2757–2765. https://doi.org/10.1242/jeb.199.12.2757

Pennycuick, C. J. (2008). Modelling the flying bird. Elsevier.

Phillips, R. A., Silk, J. R. D., Phalan, B., Catry, P., & Croxall, J. P. (2004). Seasonal sexual segregation in two Thalassarche albatross species: competitive exclusion, reproductive role specialization or foraging niche divergence? Proceedings of the Royal Society of London. Series B: Biological Sciences, 271(1545), 1283–1291. https://doi.org/10.1098/rspb.2004.2718

Pinheiro, J., Bates, D., DebRoy, S., Sarkar, D., Heisterkamp, S., Van Willigen, B., & Maintainer, R. (2017). Package ‘nlme.’ Linear and Nonlinear Mixed Effects Models, Version, 3(1). https://svn.r-project.org/R-packages/trunk/nlme/

Quintana, F., Wilson, R., Dell’Arciprete, P., Shepard, E., & Laich, A. G. (2011). Women from Venus, men from Mars: inter-sex foraging differences in the imperial cormorant Phalacrocorax atriceps a colonial seabird. Oikos, 120(3), 350–358. https://doi.org/10.1111/j.1600-0706.2010.18387.x

R Core Team. (2020). R: A language and environment for statistical computing. http://www.r-project.org/index.html

Rosén, M., Spedding, G. R., & Hedenström, A. (2007). Wake structure and wingbeat kinematics of a house-martin Delichon urbica. Journal of the Royal Society Interface, 4(15), 659–668. https://doi.org/10.1098/rsif.2007.0215

Rosén, M., Spedding, G. R., & Hedenstrom, A. (2004). The relationship between wingbeat kinematics and vortex wake of a thrush nightingale. Journal of Experimental Biology, 207(24), 4255–4268. https://doi.org/10.1242/jeb.01283

Rosén, M., Spedding, G. R., & Hedenström, A. (2007). Wake structure and wingbeat kinematics of a house-martin Delichon urbica. Journal of The Royal Society Interface, 4(15), 659–668. https://doi.org/10.1098/rsif.2007.0215

Sato, K., Daunt, F., Watanuki, Y., Takahashi, A., & Wanless, S. (2008). A new method to quantify prey acquisition in diving seabirds using wing stroke frequency. Journal of Experimental Biology, 211(1), 58–65. https://doi.org/10.1242/jeb.009811

Schmidt-Wellenburg, C. A., Biebach, H., Daan, S., & Visser, G. H. (2007). Energy expenditure and wing beat frequency in relation to body mass in free flying Barn Swallows (Hirundo rustica). Journal of Comparative Physiology B, 177(3), 327–337. https://doi.org/10.1007/s00360-006-0132-5

Shepard, E., Wilson, R., Quintana, F., Gómez Laich, A., Liebsch, N., Albareda, D., Halsey, L., Gleiss, A., Morgan, D., Myers, A., Newman, C., & McDonald, D. (2008). Identification of animal movement patterns using tri-axial accelerometry. Endangered Species Research, 10(1), 47–60. https://doi.org/10.3354/esr00084

Shirai, M., Niizuma, Y., Tsuchiya, K., Yamamoto, M., & Oka, N. (2013). Sexual size dimorphism in Streaked Shearwaters Calonectris leucomelas. Ornithological Science, 12(1), 57–62. https://doi.org/10.2326/osj.12.57

Shyy, W., Aono, H., Chimakurthi, S. K., Trizila, P., Kang, C. K., Cesnik, C. E. S., & Liu, H. (2010). Recent progress in flapping wing aerodynamics and aeroelasticity. Progress in Aerospace Sciences, 46(7), 284–327. https://doi.org/10.1016/j.paerosci.2010.01.001

Spear, L. B., & Ainley, D. G. (1997). Flight behaviour of seabirds in relation to wind direction and wing morphology. Ibis, 139(2), 221–233. https://doi.org/10.1111/j.1474-919x.1997.tb04620.x

Spivey, R. J., & Bishop, C. M. (2013). Interpretation of body-mounted accelerometry in flying animals and estimation of biomechanical power. Journal of the Royal Society Interface, 10(87). https://doi.org/10.1098/rsif.2013.0404

Taylor, L. A., Taylor, G. K., Lambert, B., Walker, J. A., Biro, D., & Portugal, S. J. (2019). Birds invest wingbeats to keep a steady head and reap the ultimate benefits of flying together. PLOS Biology, 17(6), e3000299. https://doi.org/10.1371/journal.pbio.3000299

Tobalske, B. W., Hedrick, T. L., Dial, K. P., & Biewener, A. A. (2003). Comparative power curves in bird flight. Nature, 421(6921), 363–366. https://doi.org/10.1038/nature01284

Tobalske, B. W., & Biewener, A. A. (2008). Contractile properties of the pigeon supracoracoideus during different modes of flight. Journal of Experimental Biology, 211(2), 170–179. https://doi.org/10.1242/jeb.007476

Tobalske, B. W., & Dial, K. P. (1996). Flight kinematics of black-billed magpies and pigeons over a wide range of speeds. Journal of Experimental Biology, 199(2), 263–280. https://doi.org/10.1242/jeb.199.2.263

Tobalske, B. W., Peacock, W. L., & Dial, K. P. (1999). Kinematics of flap-bounding flight in the zebra finch over a wide range of speeds. Journal of Experimental Biology, 202(13), 1725–1739. https://doi.org/10.1242/jeb.202.13.1725

Tobalske, B. W, Warrick, D. R., Clark, C. J., Powers, D. R., Hedrick, T. L., Hyder, G. A., & Biewener, A. A. (2007). Three-dimensional kinematics of hummingbird flight. Journal of Experimental Biology, 210(13), 2368–2382. https://doi.org/10.1242/jeb.005686

Torre-Bueno, J. R., & Larochelle, J. (1978). The metabolic cost of flight in unrestrained birds. Journal of Experimental Biology, 75(September 1978), 223–229. https://doi.org/10.1242/jeb.75.1.223

Tucker, B. Y. V. A. (1968). Respiratory Exchange and Evaporative Water Loss in the Flying Budgerigar. Journal of Experimental Biology, 48(1), 67–87. https://doi.org/10.1242/jeb.48.1.67

Usherwood, J. R., Stavrou, M., Lowe, J. C., Roskilly, K., & Wilson, A. M. (2011). Flying in a flock comes at a cost in pigeons. Nature, 474(7352), 494–497. https://doi.org/10.1038/nature10164

Van Walsum, T. A., Perna, A., Bishop, C. M., Murn, C. P., Collins, P. M., Wilson, R. P., & Halsey, L. G. (2020). Exploring the relationship between flapping behaviour and accelerometer signal during ascending flight, and a new approach to calibration. Ibis, 162(1), 13–26. https://doi.org/10.1111/ibi.12710

Wang, Y., Yin, Y., Ge, S., Li, M., Zhang, Q., Li, J., Wu, Y., Li, D., & Dudley, R. (2019). Limits to load-lifting performance in a passerine bird: The effects of intraspecific variation in morphological and kinematic parameters. PeerJ, 2019(11), 1–12. https://doi.org/10.7717/peerj.8048

Ward, S., Möller, U., Rayner, J. M. V., Jackson, D. M., Bilo, D., Nachtigall, W., & Speakman, J. R. (2001). Metabolic power, mechanical power and efficiency during wind tunnel flight by the European starling Sturnus vulgaris. Journal of Experimental Biology, 204(19), 3311–3322. https://doi.org/10.1242/jeb.204.19.3311

Warham, J. (1977). Wing loadings, wing shapes, and flight capabilities of Procellariiformes. New Zealand Journal of Zoology, 4(1), 73–83. https://doi.org/10.1080/03014223.1977.9517938

Weimerskirch, H., Louzao, M., De Grissac, S., & Delord, K. (2012). Changes in wind pattern alter albatross distribution and life-history traits. Science, 335(6065), 211–214. https://doi.org/10.1126/science.1210270

Wilson, R., & Liebsch, N. (2003). Up-beat motion in swinging limbs: new insights into assessing movement in free-living aquatic vertebrates. Marine Biology, 142(3), 537–547. https://doi.org/10.1007/S00227-002-0964-9

Wilson, R. P., Börger, L., Holton, M. D., Scantlebury, D. M., Gómez-Laich, A., Quintana, F., Rosell, F., Graf, P. M., Williams, H., Gunner, R., Hopkins, L., Marks, N., Geraldi, N. R., Duarte, C. M., Scott, R., Strano, M. S., Robotka, H., Eizaguirre, C., Fahlman, A., & Shepard, E. L. C. (2020). Estimates for energy expenditure in free-living animals using acceleration proxies: A reappraisal. Journal of Animal Ecology, 89(1), 161–172. https://doi.org/10.1111/1365-2656.13040

Wilson, R. P., Pütz, K., Peters, G., Culik, B., Scolaro, J. A., Charrassin, J.-B., & Ropert-Coudert, Y. (1997). Long-term attachment of transmitting and recording devices to penguins and other seabirds. Wildlife Society Bulletin, 101–106.

Wilson, R. P., White, C. R., Quintana, F., Halsey, L. G., Liebsch, N., Martin, G. R., & Butler, P. J. (2006). Moving towards acceleration for estimates of activity-specific metabolic rate in free-living animals: The case of the cormorant. Journal of Animal Ecology, 75(5), 1081–1090. https://doi.org/10.1111/j.1365-2656.2006.01127.x

